# The effect of social structure on vocal flexibility in monk parakeets

**DOI:** 10.1101/2024.09.20.614070

**Authors:** Simeon Q. Smeele, Juan Carlos Senar, Mary Brooke McElreath, Lucy M. Aplin

**Affiliations:** Cognitive & Cultural Ecology Research Group, Max Planck Institute of Animal Behavior, Radolfzell, Germany; Department of Human Behavior, Ecology and Culture, Max Planck Institute for Evolutionary Anthropology, Leipzig, Germany; Department of Biology, University of Konstanz, Konstanz, Germany; Museu de Ciències Naturals de Barcelona, Barcelona, Spain; Department of Evolutionary Biology and Environmental Science, University of Zurich, Zurich, Switzerland; Division of Ecology and Evolution, Research School of Biology, The Australian National University, Canberra, Australia

**Author notes:** **Author for correspondence:** Simeon Q. Smeele. Department of Ecoscience, Aarhus Univerisity, Aarhus, Denmark (Simeon Q. Smeele). Joint senior author (Mary Brooke McElreath, Lucy M. Aplin). **Subject areas:** behaviour.

**Keywords:** vocal complexity, social complexity, monk parakeet, parrots, Bayesian statistics

## Abstract

The social complexity hypothesis argues that communicative complexity arises as a result of social complexity, with this occurring through mechanisms including plasticity and selection. Most research to date has focused on ultimate drivers of repertoire size, for example finding that cooperative breeding species exhibit larger repertoires. Until this date no study has focused on individual-level drivers of vocal diversity. Here, we examine social networks and vocalisations in wild colonial-nesting monk parakeets (*Myiopsitta monachus*). First, we recorded social networks for 337 individuals, relatedness for 100 individuals and matched these with 5599 vocalisations from 229 individuals over two years. Overall, we found that all individuals exhibited high contact-call diversity; however, individual-level diversity increased with age in 2020 and with number of nest mates in 2021. Call similarity was not predicted by relatedness, but individuals with stronger affiliative bonds had more dissimilar calls, suggesting an active process to sound unique among close associates. Finally, females had more diverse repertoires, producing relatively fewer contact calls across years and individuals living in larger groups had more diverse repertoires in 2021. Our results demonstrate a multi-faceted social influence on call content, diversity and repertoire diversity, exhibiting how fine-scale variation in social structure can influence expressed vocal complexity.

## Introduction

Animal species differ widely in the flexibility and complexity of their vocal communication systems. Some species exhibit relatively fixed and simple vocal repertoires, while other species share information about predator type (Seyfarth, Cheney, and Marler 1980), intent to move (Arnold and Zuberbü hler 2008), and individual (Thomsen, Balsby, and Dabelsteen 2013) or group identity (Gero, Hal Whitehead, and Rendell 2016). The evolutionary drivers of this variation in vocal complexity have been of long standing interest to researchers. Most notably, Freeberg (2006) put forward the ‘social complexity hypothesis’, which states that communicative complexity arises as a result of a need to navigate social interactions and dilemmas in larger or more complex groups. Multiple studies have since used comparative analyses to examine the correlates between vocal repertoire size or diversity and various measures of sociality in taxonomic groups ranging from primates (Bouchet, Blois-Heulin, and Lemasson 2013), bats (Knö rnschild, Fernandez, and Nagy 2020) and birds (Leighton 2017) – for a summary see Peckre, Kappeler, and Fichtel (2019). This body of work has suggested a general pattern whereby cooperatively breeding species, or species living in larger groups, have more diverse vocalisation types, often summarised as ‘larger groups have more to talk about’ (Pollard and Blumstein 2012).

Many species exhibit vocal learning, with songs and calls shaped by interactions with parents and peers, either in an early developmental period, or throughout life. In these species, individual or population level vocal repertoire size and diversity can additionally be shaped by this learning process. For instance, even without active selection, copy error and innovation can lead to the appearance of novel variants of vocalisations (Nowicki et al. 2001; Slater and Lachlan 2003; Williams et al. 2013). Cultural evolutionary models such as the ‘primary ratchet model’ argue that social structure will then affect how new behaviours spread in the population and whether they are retained over generations, with larger or better connected populations holding a more diverse distribution of variants (McElreath et al. 2018). In line with this prediction, several recent studies have found a decline in song complexity in declining populations of songbird species (Paxton et al. 2019; Crates et al. 2021). Similarly, the social complexity hypothesis predicts that within flexible communication systems, individuals in larger groups will acquire or express more complex vocalisations (Freeberg 2006). In the best explored case study, Freeberg (2006) first found that wild Carolina chickadees (*Poecile carolinensis*) interacting in larger groups had greater entropy in the note composition of the sequences they uttered. Chickadees were then caught and kept in aviaries under different group sizes. After several

weeks, birds in larger groups produced social calls with the potential for greater information content than those in small groups, suggesting that communicative complexity was a facultative response to the social context. However, within-species tests of the social complexity hypothesis remain rare. Within a species, sociality can influence vocal behaviour in a different way as well. Individuals that are more closely associated can actively converge vocally to signal group identity and groups can vocally converge to signal the willingness to fuse during foraging (Janik and Slater 2000; Bradbury and Balsby 2016; Sewall 2009; Boughman 1998; Hile and Striedter 2000; Mundinger 1970; Nowicki 1989) leading to the situation where closely associated individuals and groups spending time foraging together sound more similar.

The vast majority of previous work has focused on ultimate questions of the co-evolution of vocal diversity and social complexity using comparative analyses (Peckre, Kappeler, and Fichtel 2019). We therefore still lack tests of these hypotheses at the individual level (but see Deecke et al. (2010)). Such close examination in known individuals gives the opportunity to go beyond measures of group size to interrogate what types of social relationships and social structures are most likely to impact vocal diversity in vocal learning species. Yet vocalisations often perform multiple functions across different contexts, therefore individual-level vocal diversity and repertoires may be influenced by different types of social relationships. Social network analysis enables a detailed description of social structure that spans from the individual to the population (Farine and Hal Whitehead 2015). In particular, recent developments in multiplex networks allow for multiple types of associations (e.g., foraging, aggression and spatial proximity) to be separately recorded, with metrics computed across all social networks. The resulting social measures have been shown to be qualitatively different from aggregate scores on monoplex networks in a number of studies (Sharma, Gadagkar, and Pinter-Wollman 2022; Smith-Aguilar et al. 2019; Finn et al. 2019). From the social network multiple useful metrics can be computed. Degree is an expression for how many conspecifics an individual interacts with, betweenness an expression for how important an individual is for information flow through the network and eigenvector centrality an expression for how influential an individual is in the network. From a multiplex network versatility is a measure similar to degree, but includes connections across network types.

Here, we explore the correlation between contact call and repertoire diversity and social networks in a wild population of monk parakeets (*Myiopsitta monachus*) in Barcelona, Spain. The monk parakeet is a medium sized parrot originally from central South America that has invaded and established large populations in southern Europe since the 1960s (Postigo et al. 2019). Uniquely amongst parrots, monk parakeets build communal nests from sticks, where multiple breeding pairs or cooperatively breeding trios expand on the same nest structure. Nests exist in aggregated colonies and are maintained and roosted in year round (Bucher et al. 1990; Dawson Pell, Senar, D. Franks, et al. 2021). In European cities, monk parakeets forage in fission-fusion flocks around nesting colonies (Dawson Pell, Hatchwell, Ortega-Segalerva, and Senar 2024), with foraging flocks ranging between one and 50 or more individuals, and with contact calls and other call types being uttered regularly. Parrot species have repertoires ranging from a handful of call types (Taylor and Perrin 2005) to dozens (May 2004; Farabaugh, Brown, and Dooling 1992), and can vocally learn throughout their lives (Bradbury and Balsby 2016). Earlier work on monk parakeet vocalisations has described 11 distinct call types (Martella and Bucher 1990) but only the contact call has been extensively studied (Smith-Vidaurre, Araya-Salas, and Wright 2020; Smith-Vidaurre, Perez-Marrufo, and Wright 2021; Smeele, Senar, et al. 2023; Smeele, Tyndel, et al. 2022). Previous work has suggested that these contact calls are individually distinct (Smith-Vidaurre, Araya-Salas, and Wright 2020; Smith-Vidaurre, Perez-Marrufo, and Wright 2021; Smeele, Senar, et al. 2023) but with high within-individual variability (Smeele, Senar, et al. 2023). Unlike other studied parrot species, there is no evidence for active convergence on group-level vocal signatures (Smith-Vidaurre, Araya-Salas, and Wright 2020; Wright and Dahlin 2018). However, previous work has also found information content to be lower in the American invasive range compared to the native range (Smith-Vidaurre, Perez-Marrufo, and Wright 2021), suggesting either a decline in call complexity with lower population sizes, or a historical bottleneck.

In our study we recorded vocalisations from 229 individually-marked monk parakeets in Promenade Passeig de Llúis Companys and Parc de la Ciutadella, Barcelona, Spain for two months over a two year period. We classified vocalisations into 11 call types and measured repertoire diversity, diversity in the contact call for each individual, information content in the contact call, as well as a matrix of contact-call similarity between all individuals. We combined this with a description of the social lives of individuals, where we measured foraging associations, aggressive interactions, affiliative interactions and co-nesting associations, while controlling for relatedness. We then used these data to test three predictions based on the primary ratchet model and the social complexity hypothesis.

1) We predicted individuals more central in the social network to have more diverse contact calls and to have a dominant contact call variant with greater information content. Given the multi-faceted nature of monk parakeet social structure, we hypothesised that this centrality could occur either in the nesting or foraging network, or across both, and did not make any predictions as to which was more likely. 2) We predicted that individuals with greater unweighted degree, betweenness and eigenvector centrality to have a more diverse vocal repertoire (greater entropy), either because they have opportunities to learn new call types (acquisition), or because they might encounter more social interaction types (production). 3) Previous results from parrots have shown that individuals exhibit vocal convergence with group members and mates (Farabaugh, Brown, and Dooling 1992; Farabaugh, Linzenbold, and Dooling 1994; Salinas-Melgoza and Wright 2012), with a potentially important role of convergence in pair-bonding (Hile, Burley, et al. 2005). Despite the lack of group-level signatures in monk parakeets across parks, we predicted contact calls from closely associated individuals to be more similar across all social networks within the park. Alternatively, if contact calls are largely learnt and crystallised in the nest (perhaps functioning for kin recognition), we would expect a closer relationship between vocal similarity and relatedness (Leedale et al. 2020). Finally, if contact calls purely represent individual identity and are not socially learned, there should be no effect of social relationship.

See Table 1 for all predictions and how these were modeled.

**Table 1:** Hypotheses, models, predictions, and results. Models are abbreviated versions of the full Bayesian multilevel models specified in the supplemental materials. All variables were included as varying effects. S = contact call similarity, M = mate network, R = relatedness, F = foraging network, N = nesting network, T = tolerance network, D = contact call diversity, A = age, Se = Sex, Ch = chamber size, Tr = tree size, I = contact call information content, and De = degree.

| response variable | predictor variable | model | expectation | result |
| --- | --- | --- | --- | --- |
| contact call similarity | mate network | $S \sim M + R + F + N + T$ | more similar | none |
| | foraging network | $S \sim M + R + F + N + T$ | more similar | none |
| | nest network | $S \sim M + R + F + N + T$ | more similar | more similar |
| | tolerance network | $S \sim M + R + F + N + T$ | more similar | less similar |
| | relatedness | $S \sim M + R + T$ | more similar | none |
| contact call diversity/<br>repertoire diversity | network position | $D \sim N + A + Se + T + Tr$ | more diverse | none |
| | age | $D \sim A + N + C + S$ | less diverse | more diverse |
| | sex | $D \sim N + A + Se + C + Tr$ | none | females more diverse |
| | chamber size | $D \sim C + A + Se$ | more diverse | more diverse |
| | tree size | $D \sim N + A + Se + C + Tr$ | more diverse | none |
| contact information<br>content | degree | $I \sim De + A + Tr$ | more information | none |
| | tree size | $I \sim De + A + Tr$ | more information | none |
| | age | $I \sim De + A + Tr$ | none | none |

## Methods

### Study system

The study was conducted on wild monk parakeets (*Myiopsitta monachus*) in Parc de la Ciutadella and surrounding areas in Barcelona, Spain between 27.10.20 - 19.11.20 and 31.10.21 - 30.11.21 (55 days total, see Smeele, Senar, et al. (2023) for more details).This time frame was chosen because it does not overlap with the breeding season, juveniles are largely independent, and opportunities to observe interactions between unrelated individuals are greatest. December and January were not included because the birds are less active, with temperature and light limiting the hours available for behavioural observation.

Monk parakeets have been reported as an invasive species in Barcelona since the late 1970s (Batllori and Nos 1985). From May 2002, individuals have been regularly captured and ringed using a walk-in trap on Museu de Cie`ncies Naturals de Barcelona and captured directly on their nest as fledglings (Senar, Carrillo-Ortiz, and Arroyo 2012). As a result, approximately 50% of birds were marked during the 2020 data collection period. Before the 2021 data collection period, this was supplemented with additional intensive night trapping at nests over a one week period, resulting in approximately 60-80% of the local population being marked. Individuals were ringed with stainless steel leg-bands and plastic neck-collars with a small metal medallion displaying a unique ID (Senar, Carrillo-Ortiz, and Lluisa Arroyo 2012). All birds were ringed with special permission EPI 7/2015 (01529/1498/2015) from Direcció General del Medi Natural i Biodiversitat, Generalitat de Catalunya.

During capture birds were aged as nestling, juvenile (*<*1 year) or adult. A small blood sample was taken from either the brachial or jugular vein of each individual, stored in 98% ethanol and kept at -20^°^C. Samples were then molecularly sexed by Vetgenomics at the Universitat Auto`noma de Barcelona using the P2 and P8 primers (Griffiths et al. (1998); P2 labelled with FAM). Additional sexings were obtained from Dawson Pell, Hatchwell, Ortega-Segalerva, Dawson, et al. (2020). Relatedness was then determined for 100 individuals using 21 microsatellites that were previously described by Dawson Pell, Senar, D. Franks, et al. (2021). This analysis was done by Vetgenomics (for details see supplemental materials).

### Social data collection

To quantify foraging associations, we walked random routes through Parc de la Ciutadella, where the starting point and route were randomised for each session. Data was collected for at least six hours of data collection between sunrise and sunset each day. Effort was distributed randomly throughout the day. When we encountered a single individual or group foraging or perched in any location other than a nesting tree we recorded the identity of all marked individuals. If birds were being actively provisioned by people or foraging on bread or rice we did not include the recording, because such aggregations might not represent active social association choices. We then conducted ad libitum observations in groups of displacements, fights, allopreening and close tolerance until all individuals had left or 20 minutes had passed (for details of behavioural definitions see supplemental materials).

To quantify nesting associations, we mapped the location of all nesting structures in both years in in Parc de la Ciutadella, Promenade Passeig de Llúis Companys and Zoo de Barcelona (see supplemental materials Figures 7 and 8). For each structure we assigned a tree ID, nest ID and entry ID. Each entry was considered indicative of a single nest, even though nesting chambers can have multiple entries, a single entry usually only leads to one chamber. Once nest locations were mapped, we monitored all nest entries opportunistically throughout the field season, or until we had at least three sightings of at least two individuals for a nest entry (for details see supplemental materials). Each individual was then assigned to the nest entry where it was observed most frequently. The nest network consisted of the physical distance between nesting locations for all individuals. Observations of social interactions at nest sites (e.g., allopreening) were also recorded opportunistically at nest sites.

### Social network analysis

From the observations away and at the nests (see supplemental materials for details), we generated five social networks for each data collection period. (1) The *nest network* consisted of the distance between nest locations for all individuals (0-966 m), where two nests in the same tree would be scored as 0. Values were normalised (0-1) before analysis. In addition, we recorded *tree group size* for each nesting tree, with this calculated as the number of conspecifics observed using the same nesting tree. (2) The *mate network* consisted of a binary network where all individuals sharing the same nest chamber shared an edge (for definitions of terms see Table 2). Preliminary analysis suggested that the vast majority of allopreening interactions were within mated pairs, and so observations of individuals allopreening were also incorporated into this network. In addition to this, we also recorded *chamber group size*, measured as the number of individuals assigned to a given nest entry. (3) The *foraging network* was generated from foraging observations using a *gambit of the group* approach (Hal Whitehead 2008; Croft, James, and Krause 2008; Daniel W Franks, Ruxton, and James 2010), where two individuals were given an edge if observed foraging in the same group. Edges were then scaled between 0 (never observed foraging together) to 1 (always observed foraging together) using the simple ratio index (Hubalek 1982; Sailer and Gaulin 1984; HAL Whitehead 1999; Farine and Hal Whitehead 2015). (4) The *aggression network* was constructed from observations of displacements or fights observed either during foraging transects or at the nest. Given the relatively scarce data, we collated this data into a undirected binary network, where any two individuals that were involved in an aggressive interaction were scored as one and all other zero. Finally, the (5) *affiliative network* was similar, but constructed using all observations of close tolerance behaviour and allopreening.

**Table 2:** Glossary. Definitions are based on Wey et al. (2008) and Powell (2008).

| Term | Definition |
| --- | --- |
| node | an individuals in the network |
| edge | the connection between individuals in the network, e.g., if they share a nest |
| edge weight | the strenght of the edge, e.g. based on the fraction of times individuals were observed together vs alone |
| degree | the number of edges attached to a node |
| eigenvector centrality | a measure of influence of a node in the network |
| betweenness centrality | the number of shortest paths between dyads on which the focal individual lies |
| multiplex network | a network consisting of multiple other networks where nodes present in multiple networks are connected |
| nest chamber size | number of individuals living within the same nest chamber, ranging from 1-4 |
| tree size | number of individuals living within the same tree, ranging from 1-20 |
| DAG | directed acyclic graph, representation of variables and their causal paths |
| back-door path | a non-causal path between two variables, represented by an arrow pointing towards the explanatory variable |
| indirect path | a causal path where one variable affects a second and the second affects a third variable, whereby an effect of the first on the third is observed |
| fork | a variable that has an effect on two other variables and can introduce a spurious correlation between the two |
| collider | a variable where two arrows enter, when included in the model this opens up a back-door path |

**Table 3:** Total effect for all predictor variables in the models of contact call similarity, contact call diversity, contact call information content, and repertoire entropy. Results include the average and 89% posterior interval of each effect.

| Response variable | Year | Predictor variable | Average effect | Posterior interval |
| --- | --- | --- | --- | --- |
| Contact call similarity | 2020 | Foraging network | 0.19 | -0.51; 0.90 |
| Contact call similarity | 2020 | Mate network | 0.03 | -0.41; 0.49 |
| Contact call similarity | 2020 | Spatial network | 0.44 | 0.03; 0.84 |
| Contact call similarity | 2020 | Same cluster | 0.10 | -0.20; 0.51 |
| Contact call similarity | 2020 | Aggressive network | 0.25 | -0.46; 0.95 |
| Contact call similarity | 2020 | Tolerance network | -0.19 | -1.31; 0.93 |
| Contact call similarity | 2020 | Relatedness | 0.02 | -0.37; 0.42 |
| Contact call similarity | 2021 | Foraging network | -0.06 | -0.65; 0.49 |
| Contact call similarity | 2021 | Mate network | -0.03 | -0.24; 0.15 |
| Contact call similarity | 2021 | Spatial network | -0.07 | -0.24; 0.09 |
| Contact call similarity | 2021 | Same cluster | -0.01 | -0.14; 0.11 |
| Contact call similarity | 2021 | Aggressive network | 0.19 | -0.30; 0.67 |
| Contact call similarity | 2021 | Tolerance network | 1.12 | 0.19; 2.06 |
| Contact call similarity | 2021 | Relatedness | -0.13 | -0.33; 0.06 |
| Contact call diversity | 2020 | Age | -0.21 | -0.45; 0.02 |
| Contact call diversity | 2020 | Unweighted degree | 0.09 | -0.25; 0.42 |
| Contact call diversity | 2020 | Eigenvector centrality | 0.01 | -0.32; 0.35 |
| Contact call diversity | 2020 | Betweenness | -0.20 | -0.55; 0.16 |
| Contact call diversity | 2020 | Degree versatility | 0.10 | -0.20; 0.40 |
| Contact call diversity | 2020 | Sex | -0.39 | -0.92; 0.02 |
| Contact call diversity | 2020 | Chamber size | -0.15 | -0.46; 0.17 |
| Contact call diversity | 2020 | Tree size | -0.20 | -0.55; 0.15 |
| Contact call diversity | 2021 | Age | 0.10 | -0.09; 0.28 |
| Contact call diversity | 2021 | Unweighted degree | -0.07 | -0.35; 0.20 |
| Contact call diversity | 2021 | Eigenvector centrality | -0.13 | -0.41; 0.16 |
| Contact call diversity | 2021 | Betweenness | 0.05 | -0.18; 0.29 |
| Contact call diversity | 2021 | Degree versatility | -0.04 | -0.31; 0.23 |
| Contact call diversity | 2021 | Sex | -0.11 | -0.43; 0.12 |
| Contact call diversity | 2021 | Chamber size | 0.20 | -0.02; 0.42 |
| Contact call diversity | 2021 | Tree size | 0.10 | -0.12; 0.32 |
| Information content | 2020 | Unweighted degree | 0.00 | -0.07; 0.07 |
| Information content | 2020 | Tree size | 0.00 | -0.34; 0.31 |
| Information content | 2020 | Age | 0.03 | -0.00; 0.07 |
| Information content | 2021 | Unweighted degree | 0.00 | -0.03; 0.03 |
| Information content | 2021 | Tree size | -0.01 | -0.05; 0.03 |
| Information content | 2021 | Age | -0.01 | -0.04; 0.01 |
| Repertoire entropy | 2020 | Age | -0.03 | -0.18; 0.11 |
| Repertoire entropy | 2020 | Unweighted degree | 0.09 | -0.22; 0.38 |
| Repertoire entropy | 2020 | Eigenvector centrality | 0.09 | -0.22; 0.39 |
| Repertoire entropy | 2020 | Betweenness | 0.03 | -0.44; 0.46 |
| Repertoire entropy | 2020 | Degree versatility | 0.17 | -0.03; 0.36 |
| Repertoire entropy | 2020 | Sex | 0.37 | 0.07; 0.64 |
| Repertoire entropy | 2020 | Chamber size | 0.03 | -0.15; 0.22 |
| Repertoire entropy | 2020 | Tree size | 0.39 | -0.14; 0.83 |
| Repertoire entropy | 2021 | Age | -0.01 | -0.18; 0.16 |
| Repertoire entropy | 2021 | Unweighted degree | -0.17 | -0.36; 0.02 |
| Repertoire entropy | 2021 | Eigenvector centrality | -0.20 | -0.44; 0.04 |
| Repertoire entropy | 2021 | Betweenness | -0.11 | -0.26; 0.05 |
| Repertoire entropy | 2021 | Degree versatility | -0.10 | -0.28; 0.09 |
| Repertoire entropy | 2021 | Sex | 0.20 | -0.06; 0.52 |
| Repertoire entropy | 2021 | Chamber size | -0.10 | -0.26; 0.07 |
| Repertoire entropy | 2021 | Tree size | 0.24 | 0.05; 0.44 |

We constructed a *multiplex network* that included four networks: (1) the *foraging network*, (2) the *mate network*, (3) the *aggression network* and (4) the *affiliative network*. The *nesting network* was not included due to its close similarity with the *mate network*.

From the *foraging network*, we computed the social metrics of unweighted degree, betweeness and eigenvector centrality for each individual using the packages *sna* (Butts 2022) and *igraph* (Csardi and Nepusz 2006). These metrics respectively measure for each individual: a) the number of foraging associates, b) how much they move between groups, and c) the number of foraging associates of their associates (a measure of indirect centrality). From the *multiplex network*, we calculated *degree versatility*, using a custom function that summed the unweighted degree for each individual across each networks. Here we choose unweighted degree as our measure because we were interested in how many conspecifics across all networks an individual had contact with and could potentially learn from.

### Vocalisations

We used the vocalisations previously processed for Smeele, Senar, et al. (2023). This data set consisted of 5599 vocalisations (3242 vocalisations total in 2020, with an average of 21 vocalisations per individual, numbers ranging between one and 190 vocalisations; 2357 vocalisations total in 2021, with an average of 19 vocalisations per individual, numbers ranging between one and 168 vocalisations) classified into 11 call types from 229 individually-marked monk parakeets (164 individuals in 2020, 121 individuals in 2021, with 50 individuals overlapping between years).

### Contact call similarity

To test the third prediction that more closely associated individuals have more similar contact calls we first modeled the acoustic distance between contact calls from different individuals. We used a Bayesian model with the following structure:

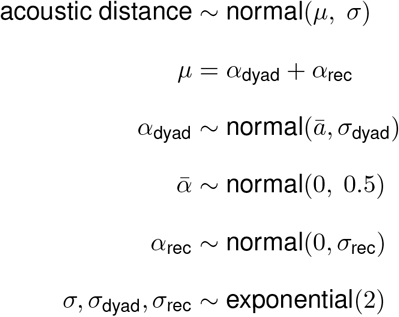

The parameter *α*_rec_ was used to control for repeated comparison of recordings. The parameter *α*_dyad_ represents the average acoustic distance between the dyads of individuals and was used in further models. We exported the mean and standard deviation to propagate uncertainty.

To model which variables could influence the similarity of contact calls between individuals, we visualised all hy-pothesised causal relationships (based on previous literature) in a directed acyclic graph (DAG, Figure 1a). Based on this, we created multilevel Bayesian models for the foraging, mate, nesting, aggressive, and tolerance networks, as well as for a matrix of pairwise genetic relatedness (see supplemental materials for details).

**Figure 1:**
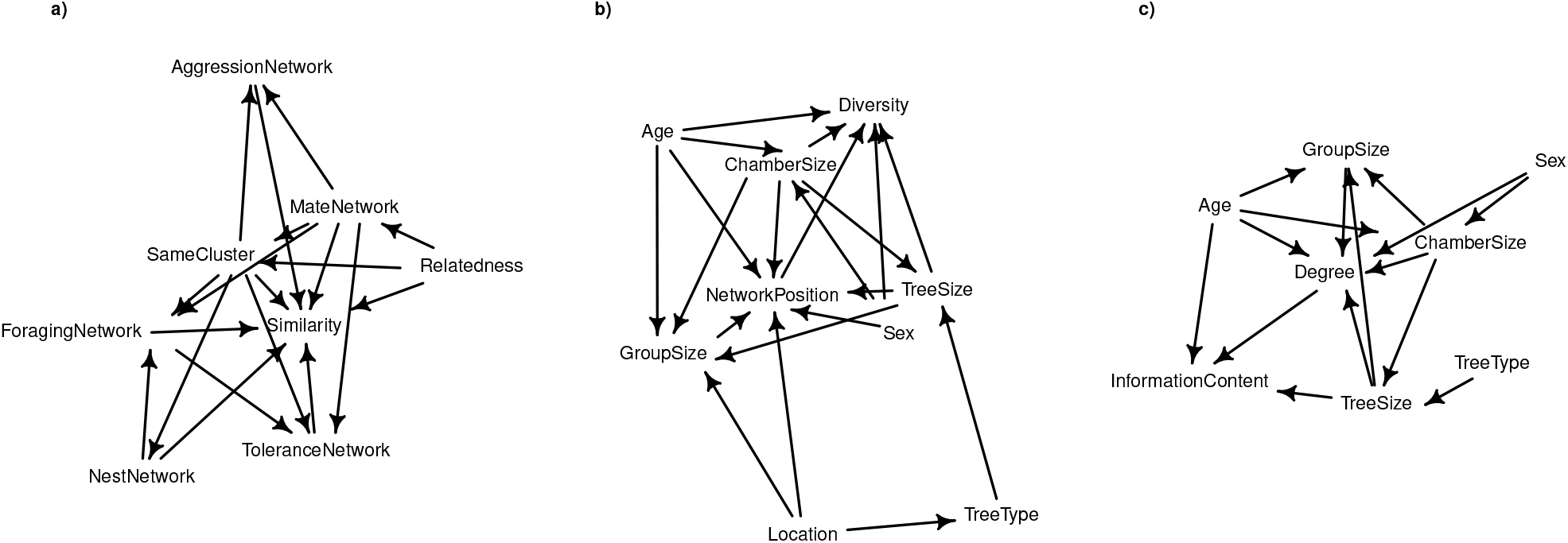
Directed acyclic graph of variables potentially influencing a) contact call similarity, b) contact call diversity and c) contact call information content. All the variables derived from the social networks are included under the term ‘NetworkPosition’. Arrows represent assumed causal effects.

### Contact call diversity

Contact calls form a near continuous set of variants within individuals (Smeele, Senar, et al. 2023). Some individuals have a more dominant variant, but these variants are not well defined. To test the first part of the first prediction, that more central individuals have more diverse contact calls, we first quantified contact call diversity by measuring the acoustic distance between contact calls from the same individual with dynamic time warping (see Smeele, Senar, et al. (2023) for details). We then used a Bayesian model with the following structure to obtain the average acoustic distance within individuals, independent of sample size and recording:

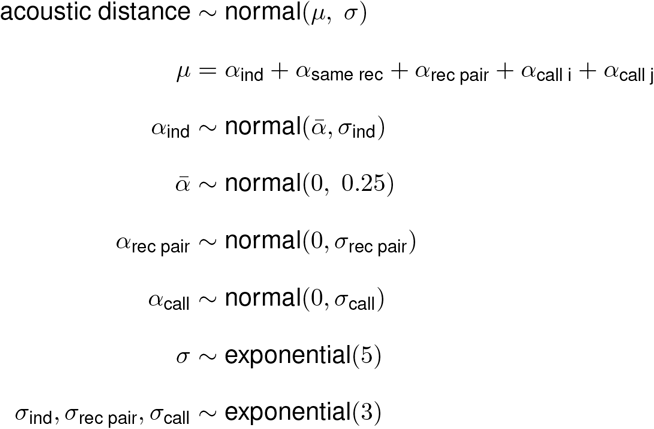

The *α*_ind_ represents the average acoustic distance between calls from the same individual and was used as the diversity measure for further analysis. Individuals that produce a larger diversity of contact calls will have a greater *α*_ind_, since the acoustic distance between dissimilar calls is greater. The parameters *α*_same rec_, *α*_rec pair_ and *α*_call_ were included to control for the fact that calls from the same recording might be more similar, calls coming from certain pairs of recording might be more similar and certain calls might be more or less typical. To propagate uncertainty about the average acoustic distance within individuals we exported both the mean and standard deviation of the posterior distribution of *α*_same rec_ for each individual.

To model which variables could potentially influence the diversity of contact calls in individuals, we first visualised all hypothesised causal relationships in a DAG (see Figure 1b). We then created Bayesian models comparing contact call diversity to age, unweighted degree, eigenvector centrality, betweenness centrality (previous three from the foraging network), degree versatility (from the multiplex network), sex, nest chamber group size and tree group size (see supplemental materials for details).

### Contact call information content

To test the second part of the first prediction that more central individuals have a dominant contact call with greater information content, we first measured the information content of contact calls for all individuals and subsetted our dataset to loud contact calls given in isolation, as defined by Smith-Vidaurre, Araya-Salas, and Wright (2020). Following Smith-Vidaurre, Perez-Marrufo, and Wright (2021) we then measured the information content by counting the number of amplitude modulation peaks by using the smoothed fundamental frequency traces from Smeele, Senar, et al. (2023) and using the function *calc*.*fm* from the R package *callsync* (**smeele2023callsync**) with the setting *min height = 30*.

To model which variables influenced the information content of contact calls in individuals, we first visualised all hypothesised causal relationships in a DAG (see Figure 1c). We then created multilevel Bayesian models comparing information content to the predictor variables of age, network position and tree group size (see supplemental materials for details).

### Repertoire diversity

To test the second prediction that more central individuals have a more diverse repertoire, we identified 11 call types across all individuals. We then quantified repertoire diversity at the individual level by calculating the entropy of each individuals repertoire:

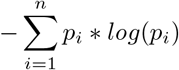

Where *p*_*i*_ is the proportion of the repertoire constituted by the i’th out of n call types in that individuals repertoire. We only included individuals with at least 30 vocalisations, to get rid of outliers created due to low sample size. This value was chosen because it ensured that either multiple recordings were included for each individual, or in a few cases, only one recording was included but this contained a warble sequence with many call types.

We modeled social and individual effects on repertoire entropy for each individual with models similar to the ones used for the contact call diversity, since we hypothesised that the explanatory variables and their causal relationships would be the same. The only prior that differed between the models was 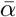,which was set to normal(1, 1).

To further test which variables influenced the composition of the repertoire, we used a modified version of the dirichlet-multinomial model proposed by Harrison et al. (2020). It models the counts of all call types for each individual with a multinomial distribution, where the probabilities for each call type occurring are drawn from a dirichlet distribution, which sums to one. The *α* parameters are the average proportions for each call type and can include fixed effects for, e.g., sex. The *β* parameters predict the increase and decline for each call type as a function of the continuous variables. We followed the same rationale as presented for the diversity analysis with respect to which variables to include for each model. For a general model definition with age as a continuous variable see below:

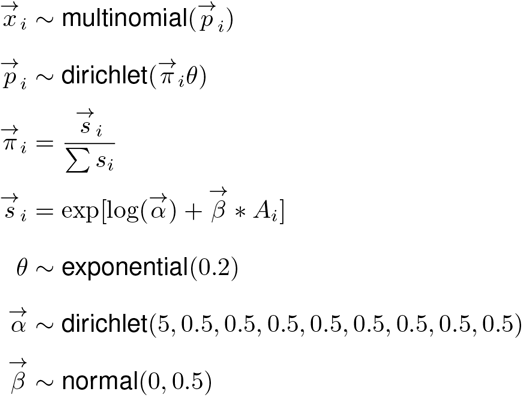

Where *θ* is a parameter controlling how variable individuals are with respect to the predicted probabilities 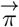.Priors were set using prior predictive simulation. We chose the prior for the 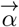 based on the knowledge that the first call type (contact call) is much more abundant than all other call types. The prior still allowed for a large range of possible values for all call types. We used the same data set as for the entropy analysis, but excluded the *frill* and *other tonal* call types, because they were too rare to be modeled reliably (see supplemental materials for details).

### Statistical analysis

All analyses were run in R (R Core Team 2021). All models were run using the package *cmdstanr* (Gabry and Č ešnovar 2021) which runs the No-U-Turn sampler (a variant of Hamiltonian Monte Carlo) based on the Stan language (Gelman, Lee, and Guo 2015). We monitored Rhat values and report if divergence occurs in the results section. For each variable we modeled the total and the direct effect by including covariates according to the back-door criterion (for details of each model see supplemental materials). To make sure we did not tailor our analysis to fit spurious results, we first analysed the data from 2021 and only after we had made the final figures replicated these with data from 2020.

## Results

### Social networks

After 23 days of data collection in October-November 2020, we had 86 individuals in the foraging network, 66 individuals in the mate network, 144 individuals in the affiliative network and 91 individuals in the aggression network (see supplemental materials Figure 6) and 175 individuals for the multiplex network. We found 44 nesting trees with 65 nests and 196 individuals (see supplemental materials Figure 7). After 31 days of data collection in November 2021 we had 129 individuals in the foraging network, 75 individuals in the mate network, 137 individuals in the tolerance network and 132 individuals in the aggression network (see Figure 2) and 164 individuals for the multiplex network. We found 34 nesting trees with 45 nests and 111 individuals (see supplemental materials Figure 8). Group size in nest chambers ranged one to four across years. Group size in nest trees ranged one to 20 across years.

**Figure 2:**
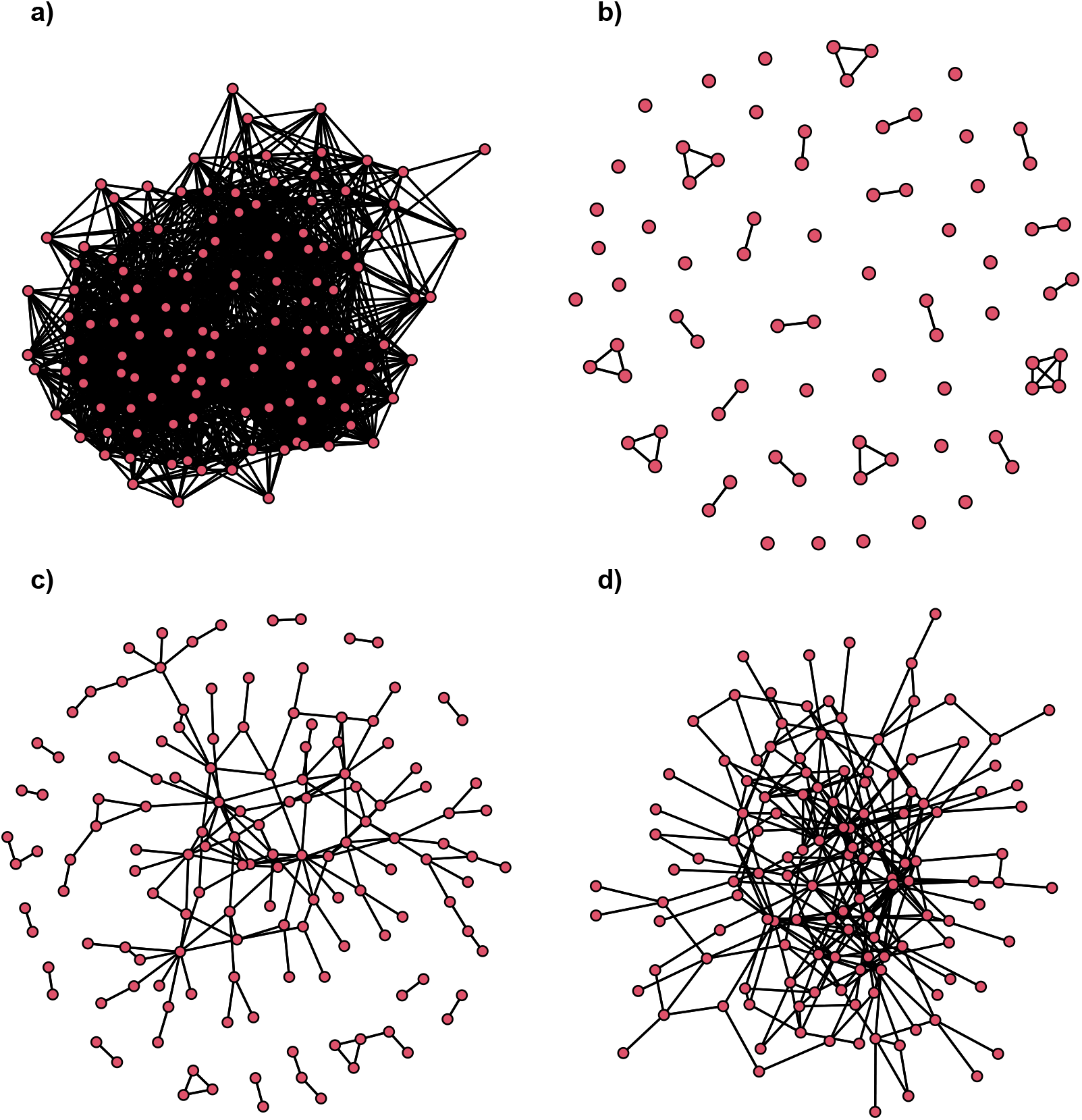
Graphs for a) foraging network, b) mate network, c) affiliative network and d) aggression network in 2021. Red nodes represent individuals, with the undirected binary edges between individuals in black. For the networks for 2020, see supplemental materials.

### Contact call similarity

Average similarity was estimated for 6551 dyads in 2020 and 5253 dyads in 2021. In general the dyadic off-set explained a modest amount of variation (*σ*_dyad_ = 0.42, 89% PI = 0.41-0.43 for 2020 and *σ*_dyad_ = 0.49, 89% PI = 0.48-0.50 for 2021).

In 2020 there was a weak total effect of the nest network on contact call similarity; individuals that nested fur-ther apart had less similar calls (see Figure 3a and -b). In 2021 this effect was absent, but there was a strong effect of the tolerance network (see Figure 3a and -c), whereby individuals that were frequently observed close together (greater edge weight) had less similar calls. None of the other variables had any effect (see supplemental materials Figure 9 and Figure 10). Contrary to our initial hypothesis, there was no effect of relatedness on call similarity.

**Figure 3:**
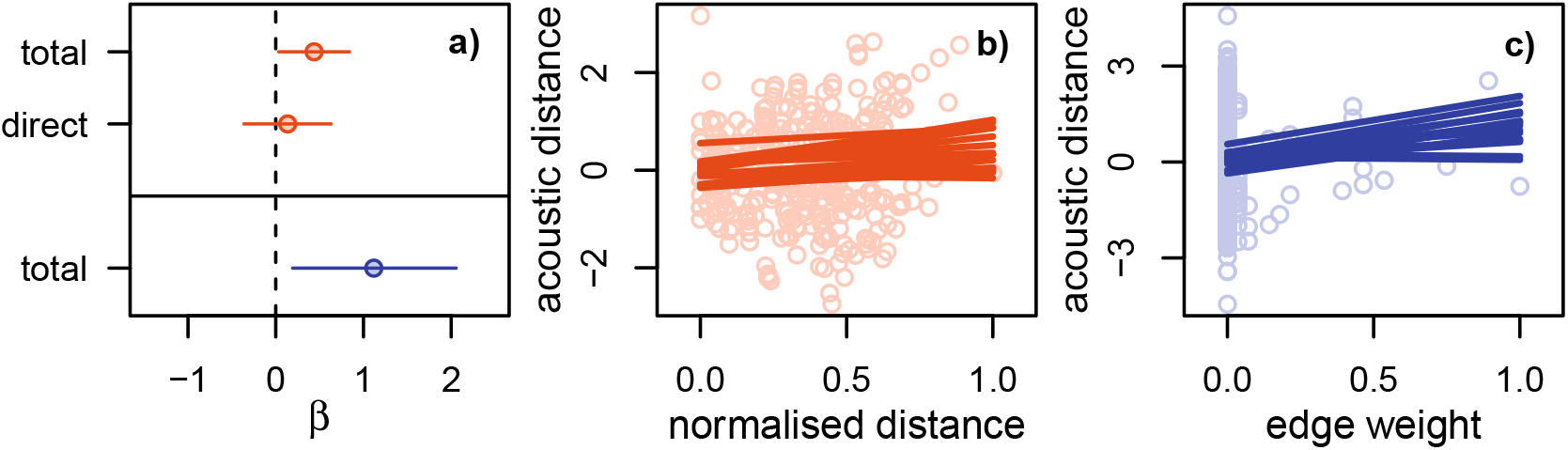
Model results for the effect of spatial (orange) and tolerance (blue) networks on contact call acoustic distance. a) Total and direct effect estimates for the spatial network in 2020 (top - orange) and total effect estimate for the tolerance network in 2021 (bottom - blue). Dots represent model average and lines represents the 89% posterior interval. b) Scatter plot of the raw data and 16 lines from the posterior distribution for the model of the direct effect of spatial network in 2020. c) Scatter plot of the raw data and 16 lines from the posterior distribution for the model of the effect of tolerance network in 2021.

### Contact call diversity

Contact call diversity was estimated with 1706 calls from 115 individuals for 2020 and with 1488 calls from 103 individuals for 2021. In general there was little variation between individuals in their contact call diversity (*σ*_ind_ = 0.04, 89% PI = 0.03-0.05 for 2020 and *σ*_ind_ = 0.05, 89% PI = 0.04-0.06 for 2021), with all individuals exhibiting a high degree of diversity. We then modelled the total and direct effect of age, sex, degree, eigenvector centrality, betweenness centrality (latter three derived from foraging network), degree versatility (multiplex network), nest chamber group size and nest tree group size on contact call diversity. We had data on age and sex for 337 individuals across both years, degree, eigenvector centrality and betweenness centrality for 90 individuals in 2020 and 128 individuals in 2021, nest chamber group size for 109 individuals in 2020 and 81 individuals in 2021 and nest group size for 161 individuals in 2020 and 99 individuals in 2021. Age and nest chamber size had a positive direct effect in 2021, but no effect in 2020 (see supplementary materials Figure 11 and Figure 12).

### Contact call information content

We measured information content (number of amplitude modulation peaks) for 664 loud contact calls from 75 individuals in 2020 and 1005 loud contact calls from 89 individuals in 2021. None of the explanatory variables had any effect on the information content of the loud contact calls (see supplemental materials) and the variation between individuals was generally also very low (mean *σ*_ind_ across models ranging between 0.02 and 0.16).

### Repertoire diversity

In order to examine repertoire diversity, we subsetted the dataset to only include individuals with at least 30 calls, leading to a dataset that included repertoires for 38 individuals in 2020 and 34 individuals in 2021. There was an effect of sex in 2020 with females having greater entropy (see Figure 4). This effect was also supported in 2021, but the 89% posterior interval overlapped zero (see supplemental materials Figure 16). The distribution analysis supported this finding that females had greater repertoire diversity. Females produced a lower proportion of contact calls in both 2020 and 2021 (see Figure 5) and overall produced other call types more often, which explains the greater entropy in the diversity analysis. Apart from sex, we found that the nest group size had a positive effect in 2021 (see Figure 4), but overlapped zero in 2020 (see supplemental materials Figure 15). For the results of the other variables see supplemental materials Figure 17.

**Figure 4:**
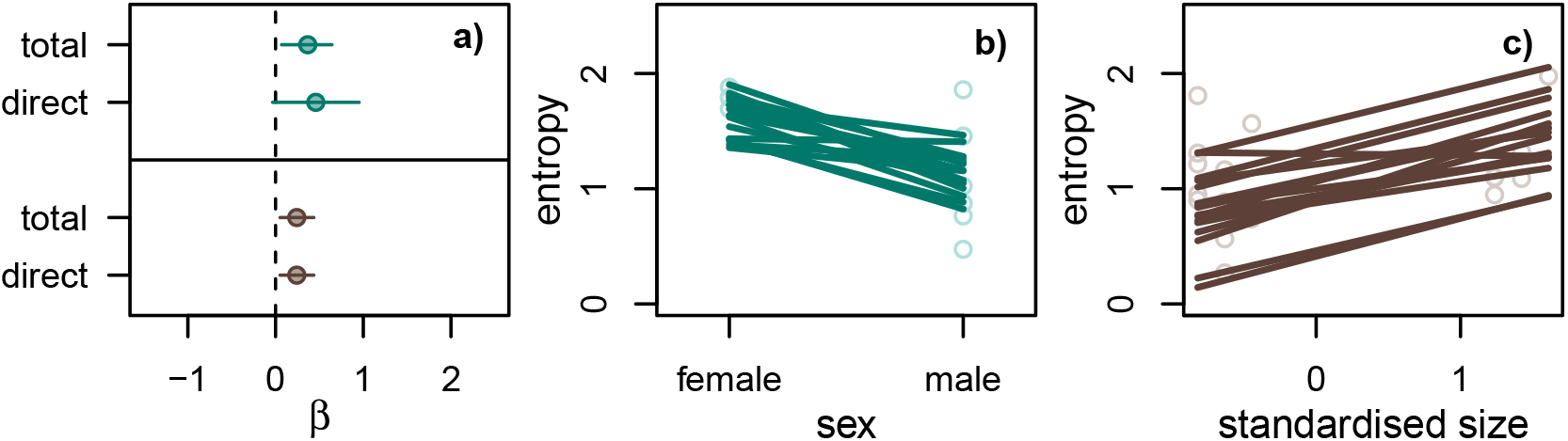
Model results for variables hypothesised to influence repertoire entropy with greatest effects. a) Total and direct effect estimates of sex in 2020 (top - green) and tree size in 2021 (bottom - brown). Dots represent model average and lines represents the 89% posterior interval. b) Scatter plot of the raw data and 16 lines from the posterior distribution of the model for the direct effect of sex in 2020. c) Scatter plot of the raw data and 16 lines from the posterior distribution of the model for the direct effect of tree size in 2021.

**Figure 5:**
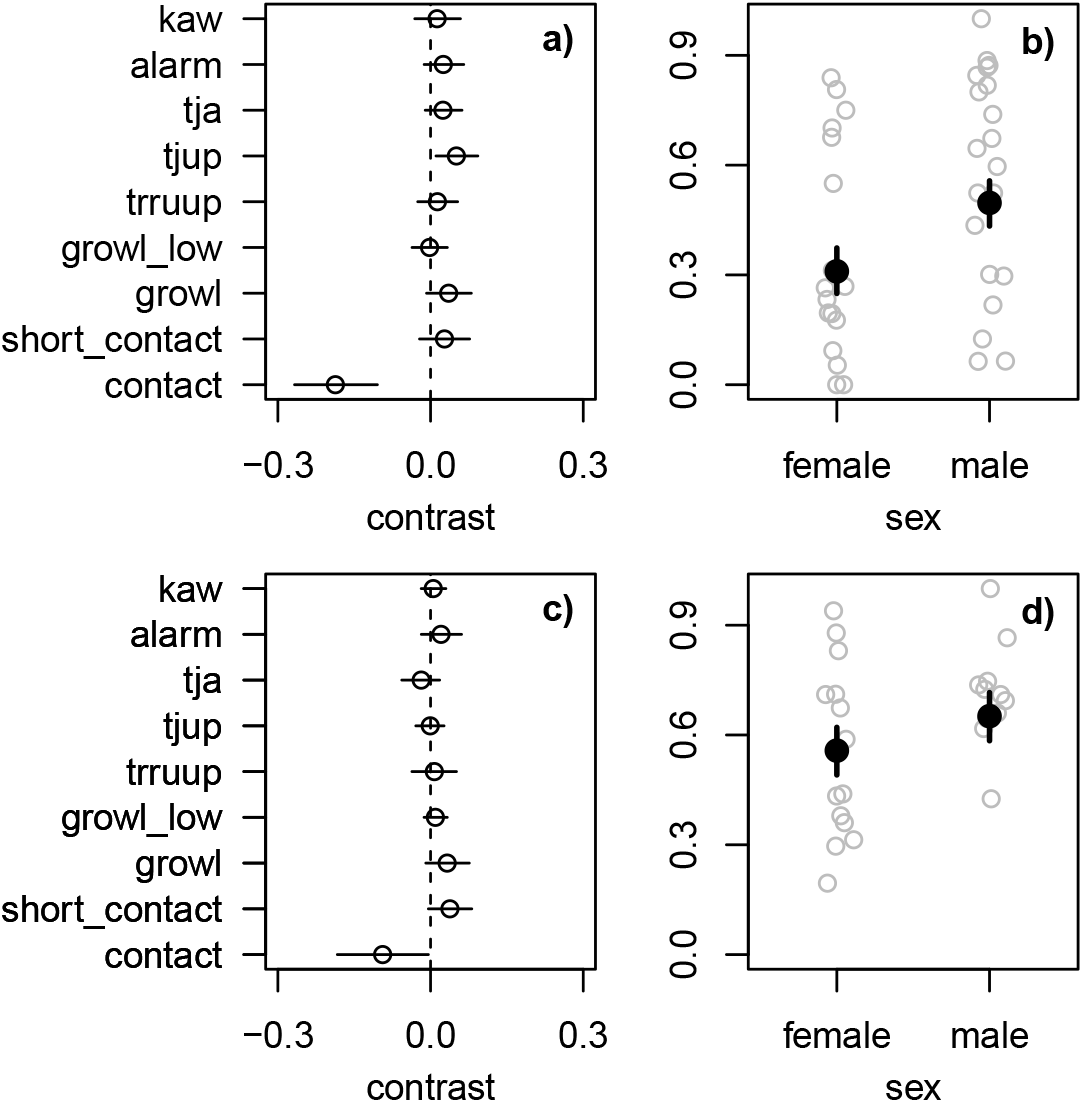
Model results for effect of sex on repertoire distribution. a) Contrast estimates between sexes (α_F™_ α_M_) for each call type in 2020. b) Scatterplot of the proportion of contact calls for females and males in 2020. Grey circles are raw data (proportion of that call type for each individual). Black dot is the mean estimate for the α parameter and the black line the 89% posterior interval. c) Contrast estimates between sexes (α_F™_ α_M_) for each call type in 2021. d) Scatterplot of the proportion of contact calls for females and males in 2021. Grey circles are raw data (proportion of that call type for each individual). Black dot is the mean estimate for the α parameter and the black line the 89% posterior interval.

**Figure 6:**
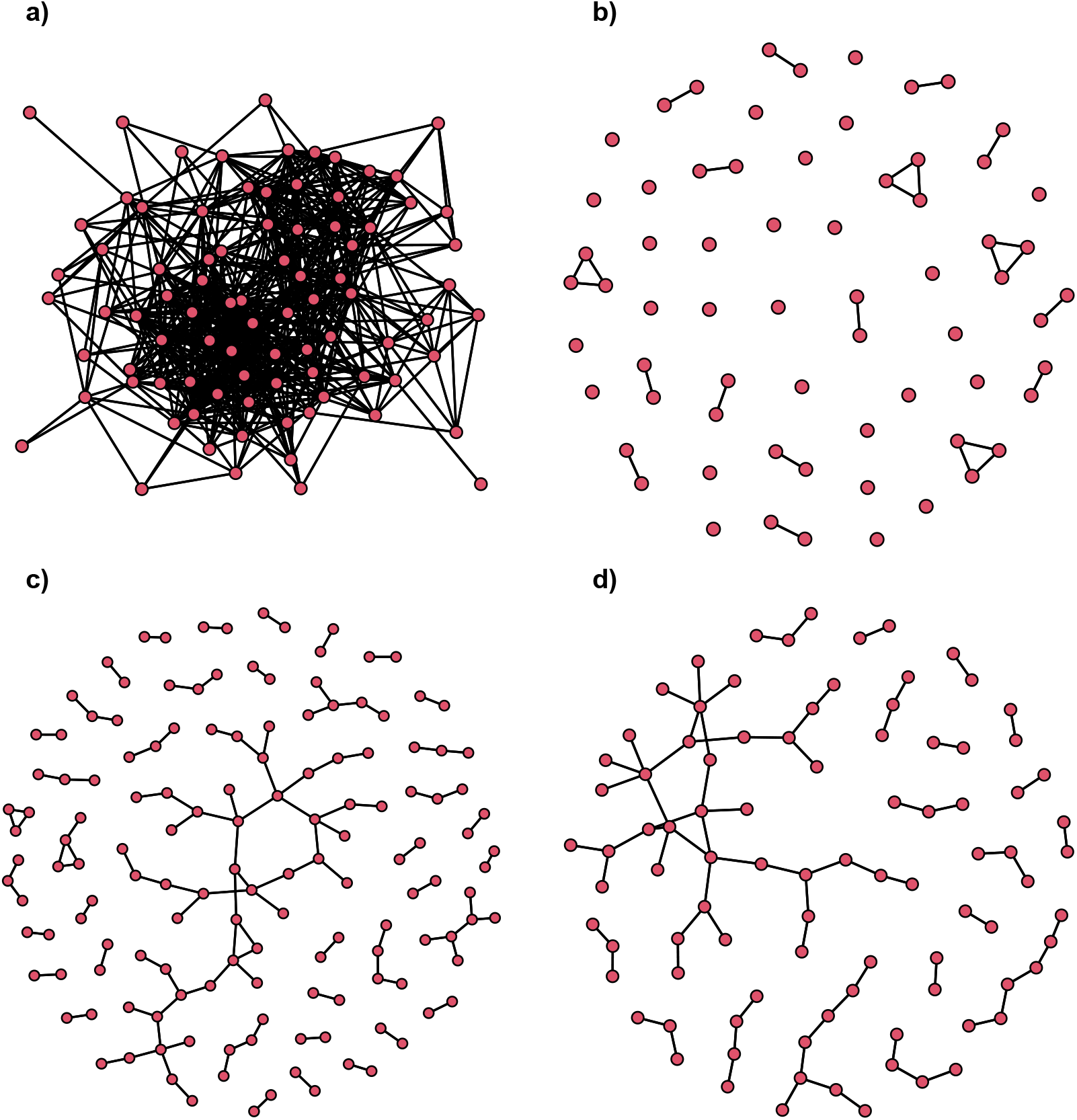
Graphs for a) foraging network, b) mate network, c) close tolerance network and d) aggression network for 2020. Red dots represent individuals. Black lines represent undirected binary edges between individuals.

**Figure 7:**
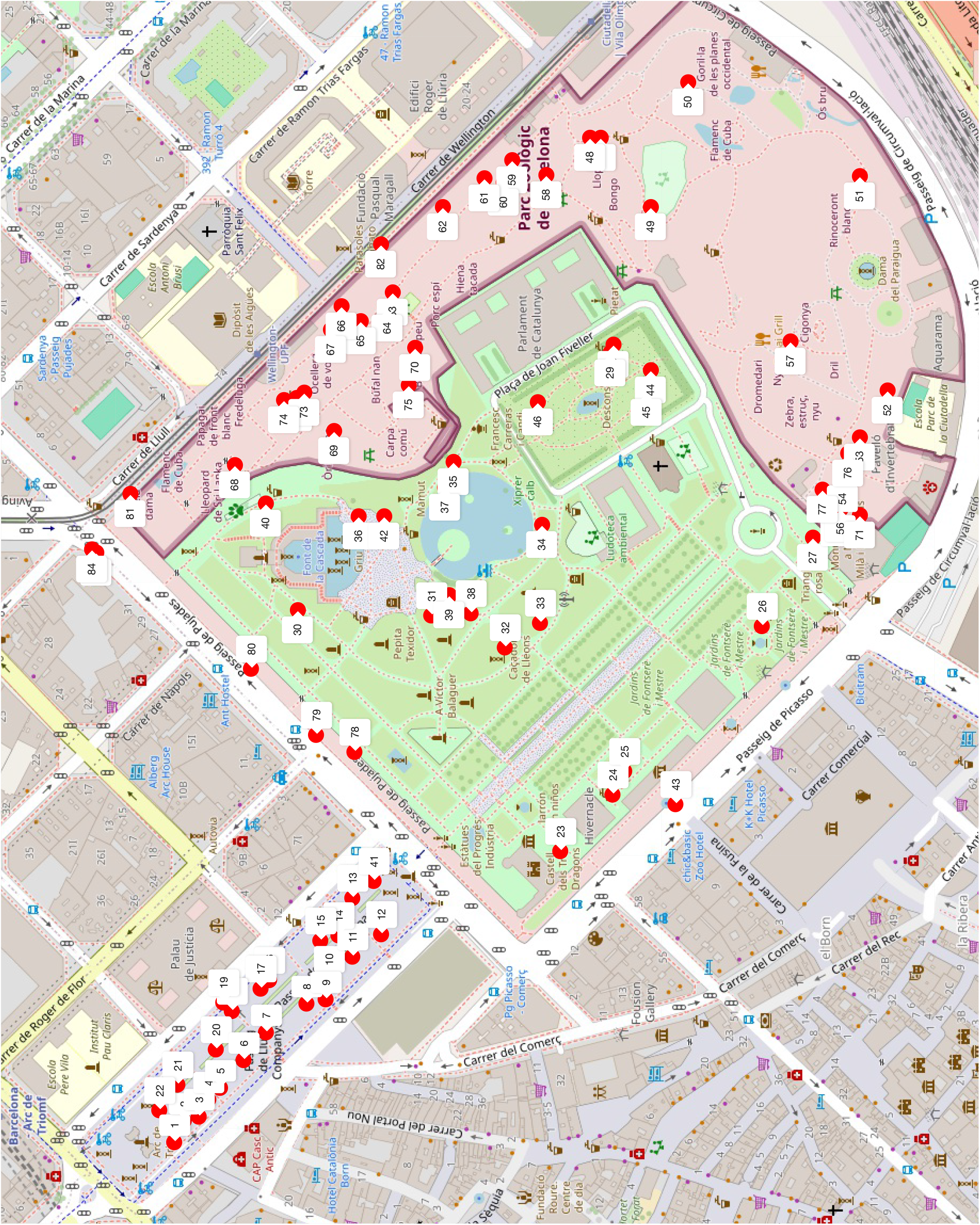
Map of Ciutadella park and surroundings with all nests from 2020. Map generated with Cheng et al. (2023)

**Figure 8:**
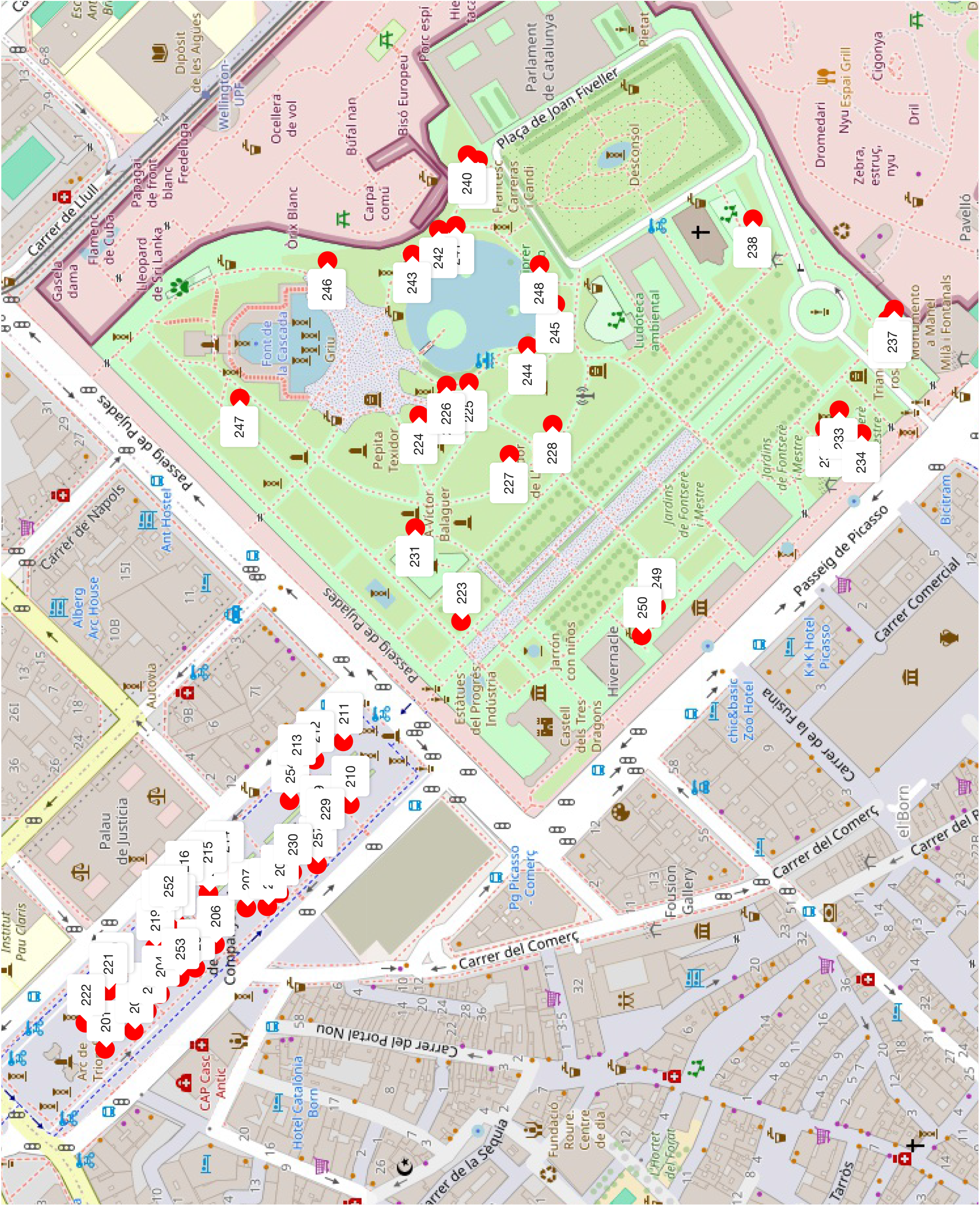
Map of Ciutadella park and surroundings with all nests from 2021. Map generated with Cheng et al. (2023)

**Figure 9:**
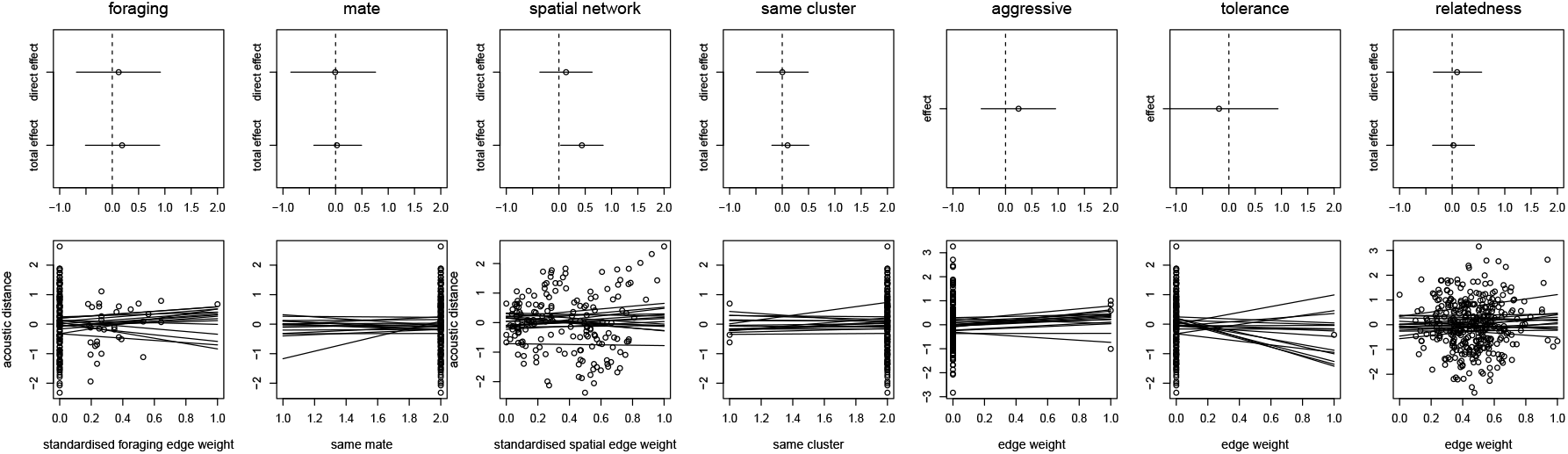
Model results for variables hypothesised to influence contact call similarity. Top: total and direct effect estimates (note that the total and direct effect for tolerance and foraging network are the same) for 2020. Dots represent model average and lines represents the 89% posterior interval. Bottom: scatter plots of the raw data and 16 lines from the posterior distribution per variable (always from the model for the total effect of that variable). Similarity (average acoustic distance between calls from the same individual) on the y-axes.

**Figure 10:**
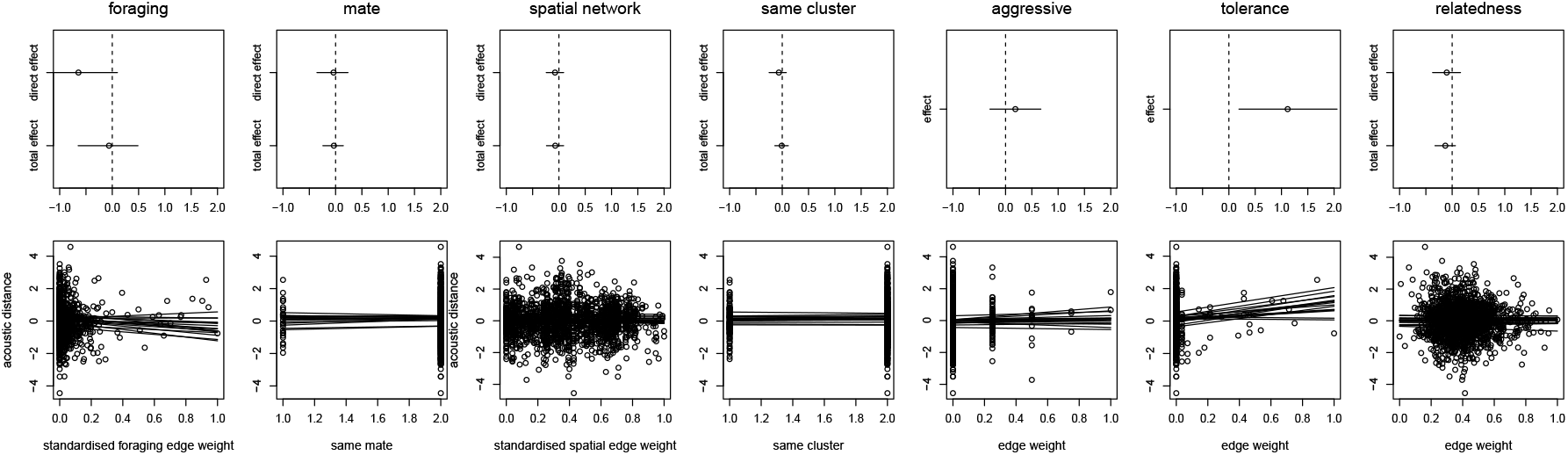
Model results for variables hypothesised to influence contact call similarity. Top: total and direct effect estimates (note that the total and direct effect for tolerance and foraging network are the same) for 2021. Dots represent model average and lines represents the 89% posterior interval. Bottom: scatter plots of the raw data and 16 lines from the posterior distribution per variable (always from the model for the total effect of that variable). Similarity (average acoustic distance between calls from the same individual) on the y-axes.

**Figure 11:**
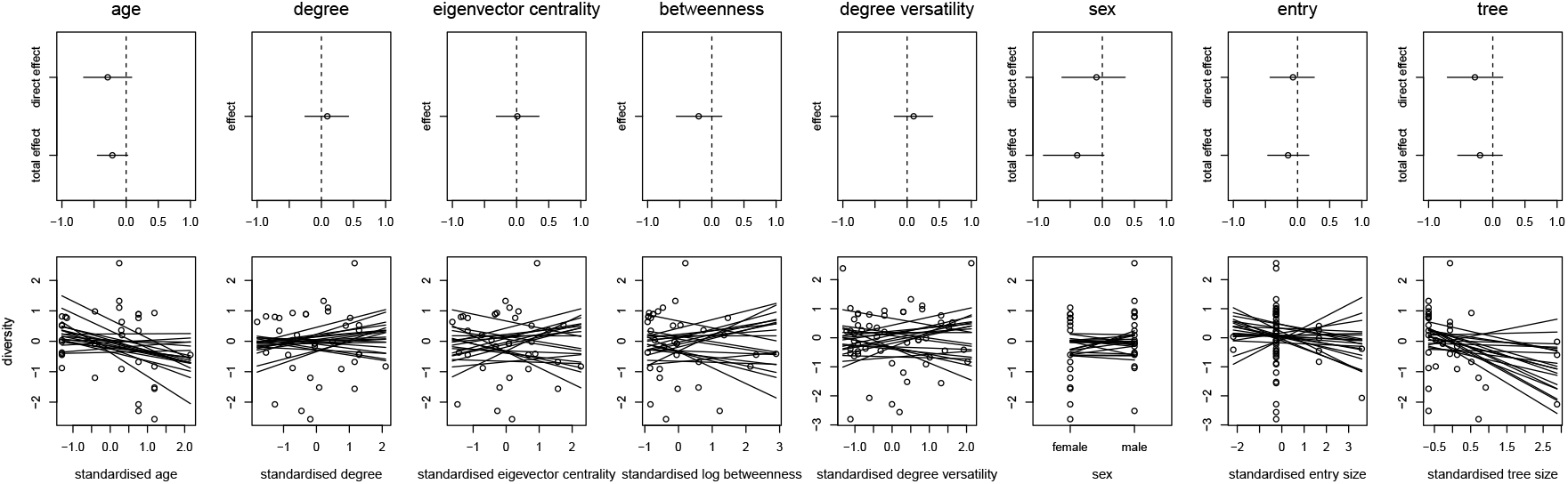
Model results for variables hypothesised to influence contact call diversity. Top: total and direct effect estimates (note that the total and direct effect for network position variables are the same) for 2020. Dots represent model average and lines represents the 89% posterior interval. Bottom: scatter plots of the raw data and 16 lines from the posterior distribution per variable (always from the model for the direct effect of that variable). Diversity (average acoustic distance between calls from the same individual) on the y-axes.

**Figure 12:**
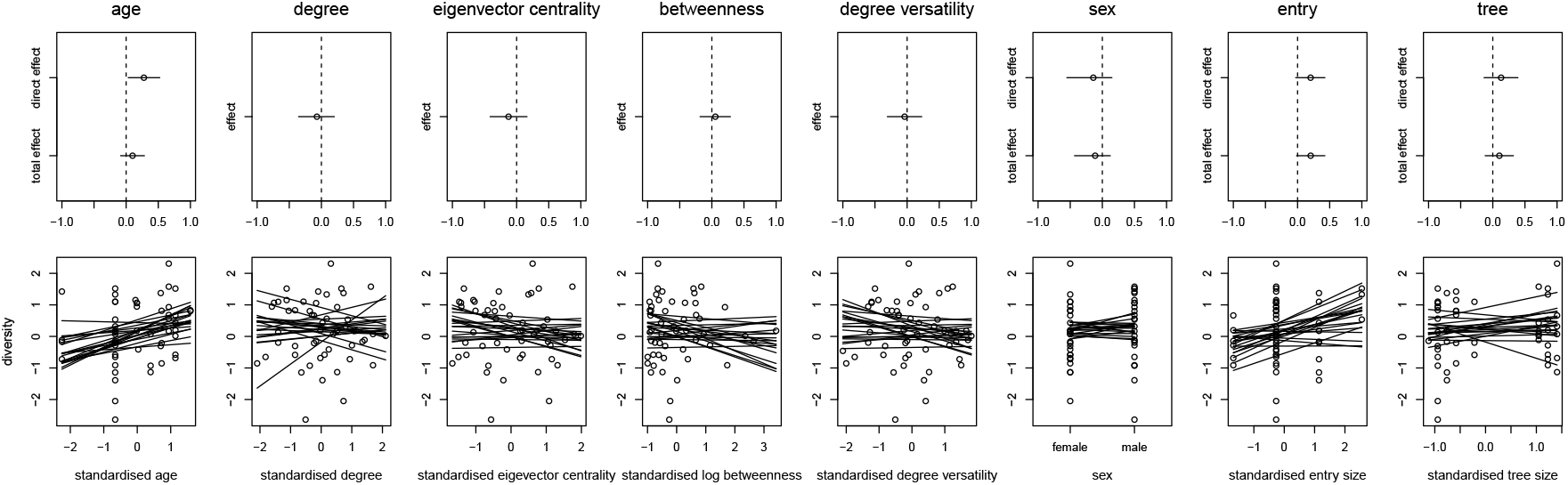
Model results for variables hypothesised to influence contact call diversity. Top: total and direct effect estimates (note that the total and direct effect for network position variables are the same) for 2021. Dots represent model average and lines represents the 89% posterior interval. Bottom: scatter plots of the raw data and 16 lines from the posterior distribution per variable (always from the model for the direct effect of that variable). Diversity (average acoustic distance between calls from the same individual) on the y-axes.

**Figure 13:**
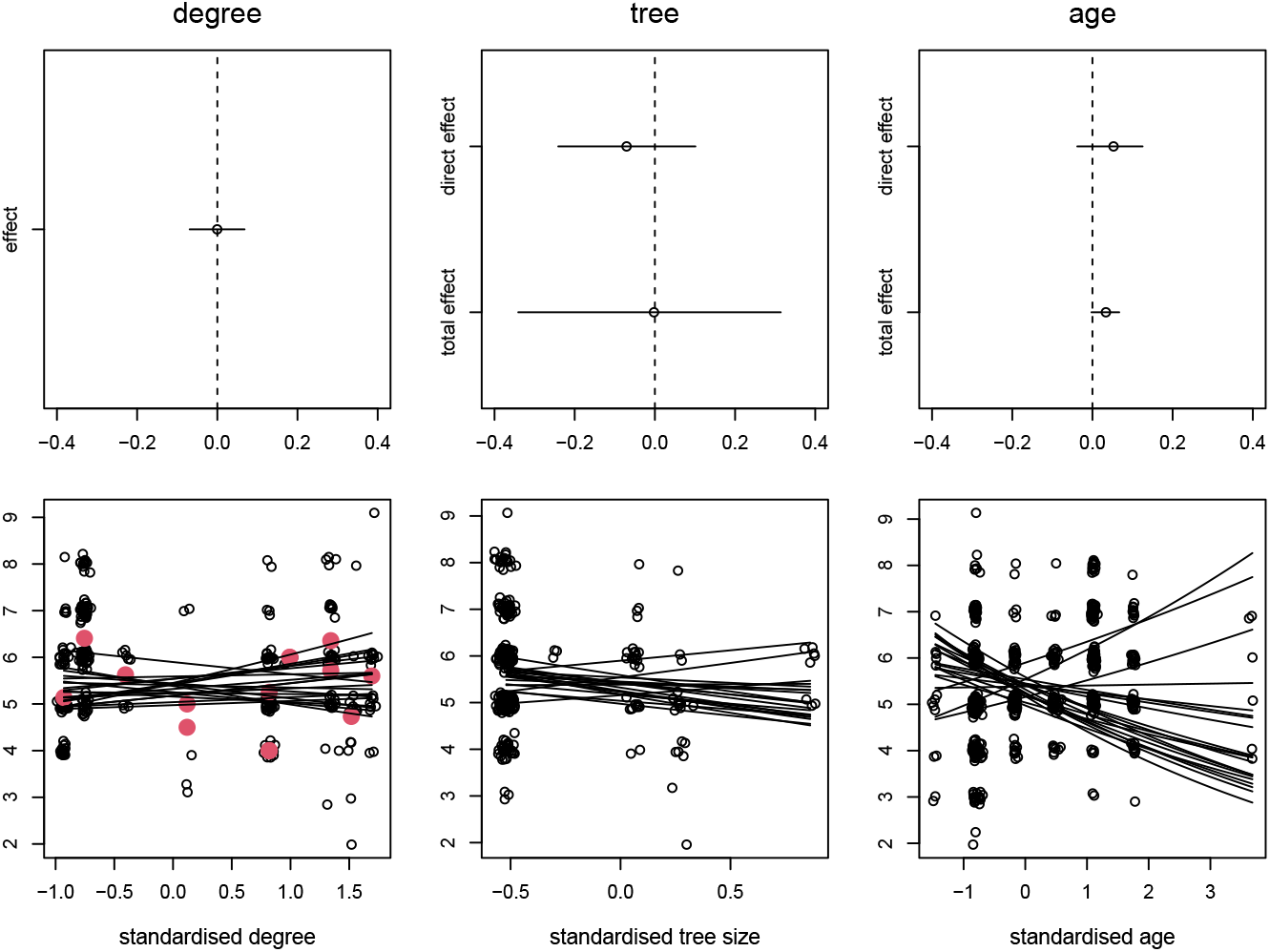
Model results for variables hypothesised to influence contact call information content for 2020. Dots represent model average and lines represents the 89% posterior interval. Bottom: scatter plots of the raw data and 16 lines from the posterior distribution per variable (always from the model for the direct effect of that variable). Red dots are individual averages.

**Figure 14:**
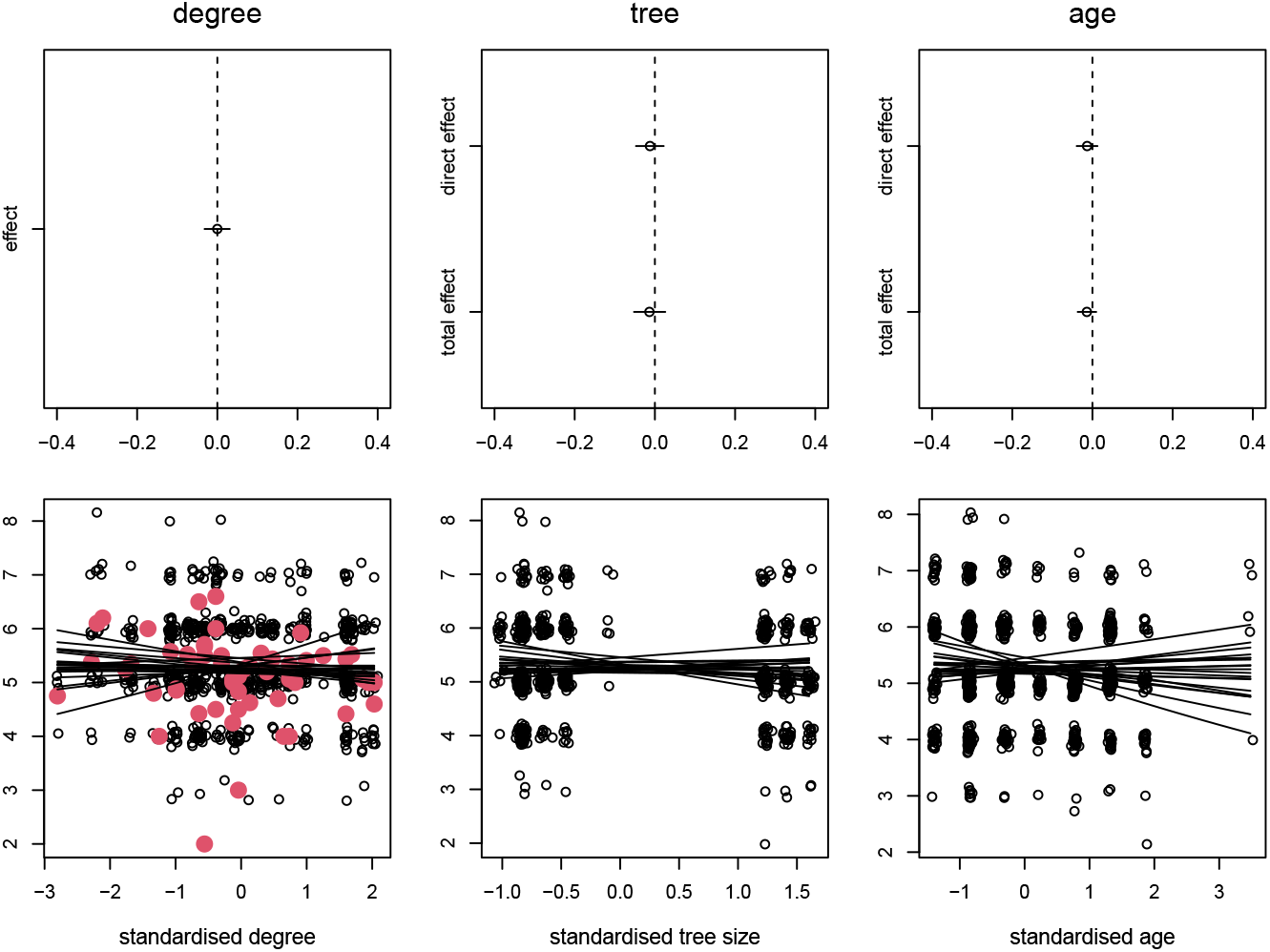
Model results for variables hypothesised to influence contact call information content for 2021. Dots represent model average and lines represents the 89% posterior interval. Bottom: scatter plots of the raw data and 16 lines from the posterior distribution per variable (always from the model for the direct effect of that variable). Red dots are individual averages.

**Figure 15:**
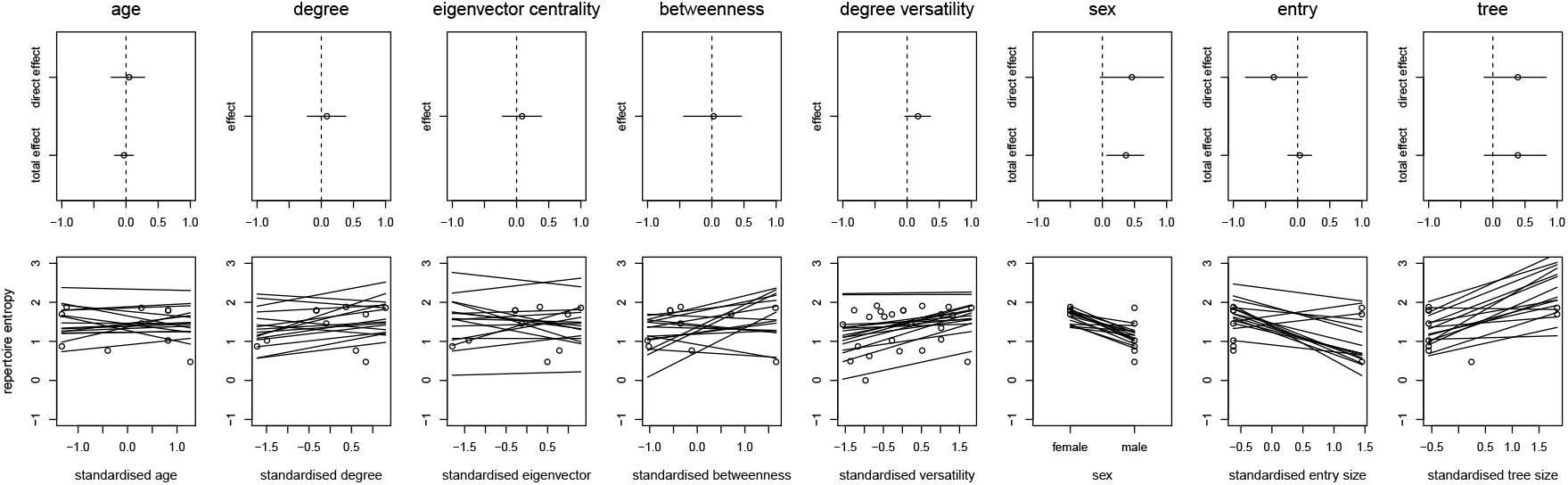
Model results for variables hypothesised to influence repertoire entropy. Top: total and direct effect estimates (note that the total and direct effect for degree versatility are the same) for 2020. Dots represent model average and lines represents the 89% posterior interval. Bottom: scatter plots of the raw data and 16 lines from the posterior distribution per variable (always from the model for the direct effect of that variable).

**Figure 16:**
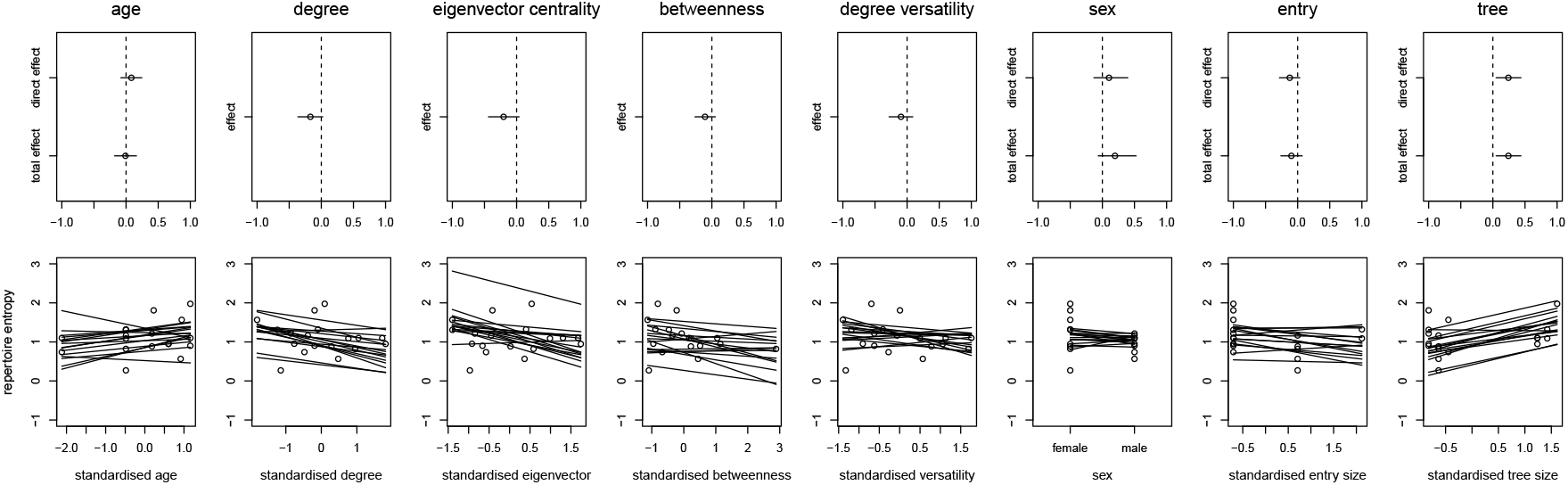
Model results for variables hypothesised to influence repertoire entropy. Top: total and direct effect estimates (note that the total and direct effect for degree versatility are the same) for 2021. Dots represent model average and lines represents the 89% posterior interval. Bottom: scatter plots of the raw data and 16 lines from the posterior distribution per variable (always from the model for the direct effect of that variable).

**Figure 17:**
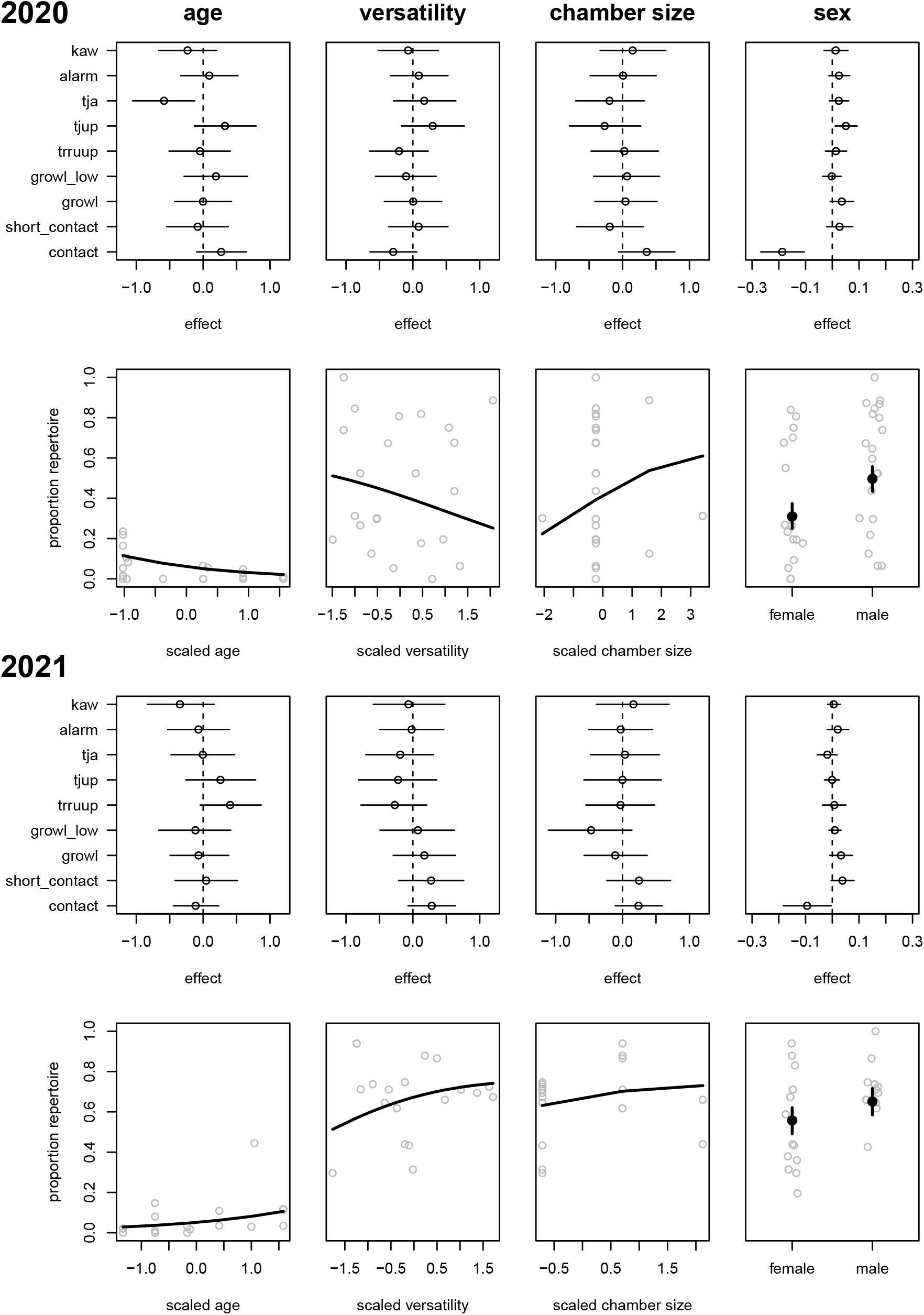
Model results for variables hypothesised to influence repertoire distributions. Top row: β parameter estimates for each call type. For sex the contrast (α_F_−α_M_) is shown. Bottom row: scatterplot for the call type with the clearest effect. Grey circles are raw data (proportion of that call type for each individual). Black line is the posterior prediction of the proportion as a function of the continuous variable. For sex the black dot is the mean estimate for the α parameter and the black line the 89% posterior interval.

## Discussion

Our study aimed to use social network approaches to test the influence of social factors on individual-level variation in vocal complexity in wild monk parakeets. In line with the social complexity hypothesis, we predicted that individuals with higher centrality in their foraging and/or nesting networks would have greater diversity and higher information content in their contact calls, and have a larger repertoire across call types. We further predicted that, if contact calls are socially influenced throughout life, then individuals should exhibit more similar vocalisations to their close associates, while if contact calls are largely learnt and crystallised in the nest, than individuals should exhibit more similar vocalisations to kin. We tested these predictions over two autumn periods, and effects varied between years, perhaps because the first field season had a lower sample size, and included fewer young individuals (mean age 2020 = 1447 days, mean age 2021 = 975) and fewer individuals living in larger nest chambers (mean tree size 2020 = 2.9 individuals, mean tree size 2021 = 3.2 individuals). However, overall, we found that most individuals had diverse contact calls, with little variation in contact call diversity or information content across individuals. Individuals had no discrete contact call variants with a dominant contact call variant, in line with the findings of Smeele, Senar, et al. (2023), but very different from, e.g., bugerigars (Farabaugh, Linzenbold, and Dooling 1994) or orange fronted conures (Bradbury, Cortopassi, et al. 2001). Contrary to the predictions, none of our social network variables predicted contact call diversity or information content, including our measure derived from the multiplex network that incorporated four types of social interactions (foraging, affiliative, foraging, nesting). Yet contact call diversity did increase with age in 2020 and with nest chamber group size in 2021. This result suggests that contact call variants continue to be acquired throughout life, and that individuals use different variants to interact with different individuals that share the same nest chamber. Individuals nesting in larger groups also had a greater repertoire diversity in 2021, although this was better predicted by the size of the nest tree rather than the nest chamber. This result is in line with the predictions of both the social complexity hypothesis and the primary ratchet model. Finally, and contrary to expectation, we didn’t find any evidence that individuals sharing a nest chamber or nest tree had more similar contact calls, although at a larger scale there was a weak effect of nesting proximity on call similarity in 2020. In addition, there was no effect of relatedness on call similarity either. Instead, there was a strong effect in 2021 for close affiliates to have greater dissimilarity in calls.

Altogether, our results suggest that nesting organisation has a much larger effect on vocal complexity in monk parakeets than foraging associations, with nesting group size at the chamber and tree level correlated with contact call diversity and repertoire diversity. Monk parakeets both roost and brood in stick nests, and work year-round to build and maintain these structures. These nests contain variable numbers of pairs and cooperative breeding groups in different nest chambers (often related: Dawson Pell, Hatchwell, Ortega-Segalerva, Dawson, et al. (2020)). Nests are valuable structures that individuals defend, for example from intruders stealing sticks, yet birds also tolerate others to build nests nearby (Dawson Pell, Senar, Ortega-Segalerva, et al. 2024). Such nesting organisation could potentially require extensive negotiation. In support of this, we observed that monk parakeets appear to use very little communication during foraging, but are very vocal around their nest (SQS personal observation). We did not measure call rate in our study, but future work could add this variable to investigate this difference more explicitly. Future studies could also examine which call types are used more frequently around the nest and how call types are used in call-response interactions, to test whether greater variability in calls allows individuals to better navigate the social interactions associated with nesting. During such a project, focus should be on recording the full repertoire of fewer individuals. One drawback of our focus on collecting social data as well, and recording as large a set of individuals as possible, is the limited amount of acoustic data for a large subset of individuals. Our modeling approach allowed us to account for the unbalanced sampling in the dirichlet-multinomial model and the contact call diversity model, but not in the model of repertoire entropy. We do not believe this introduced a bias, but results should include much less uncertainty and be less variable across years when enough acoustic data is sampled to achieve a full repertoire description for most individuals. It should also be noted that some individuals in our study were recorded in both years, resulting in some pseudo-replication across years. However, since some individuals might have shifted position in the social network and gained (or lost) vocal diversity, we opted to included data from these individuals to allow for a large enough sample size. Especially since only the lower frequency of contact calls for females was present across years, this decision should not have any effect on the overall conclusions.

Future studies could also investigate how variation in contact call diversity is distributed across time. For instance, the work by Vehrencamp et al. (2003) suggest that some diversity might be most pronounced when interacting and imitating other individuals. Also peach-fronted conures (*Eupsittula aurea*) have a stable individual call (Thomsen, Balsby, and Dabelsteen 2013), but do modify this call during interactions (Thomsen, Balsby, and Dabelsteen 2019). Similar processes could explain some of the variation observed in monk parakeets. For example, if calls are modified during vocal interactions, or dependent on the audience, variability would be high across and even within recordings. Furthermore, if individuals have a general template with relatively inflexible individuality (as found in previous studies (Smith-Vidaurre, Araya-Salas, and Wright 2020; Smeele, Senar, et al. 2023)), then we would not expect variation in the degree of contact call diversity or strong similarity between individuals, since all individuals use the same set of contact calls to interact. This notion is also supported by our findings on the information content of contact calls, where within-individual variability in how many amplitude modulation peaks were present in their contact call was high, but variability across individuals was very low.

We did find, however, that individuals tended to sound more dissimilar to their close affiliates (usually mates) than expected from the population level. We also found the surprising result that females have on average a more diverse repertoire. We did not find any effect of relatedness. Our results are not in line with a stable individual signal or a shared signal within nests or foraging groups. It is much more likely that the contact call is a flexible tool that is frequently modified. Interestingly, while most studied parrots exhibit vocal convergence in groups (Sewall, Young, and Wright 2016) and between mates (Scarl and Bradbury 2009; Hile, Plummer, and Striedter 2000), the evidence from previous studies in monk parakeets has instead found some signal of individual identity in contact calls (Smith-Vidaurre, Araya-Salas, and Wright 2020; Smeele, Senar, et al. 2023). Our results support this, but additionally suggest that the contact call can be used more flexibly than previously assumed.

To better understand the learning and usage learning of call types, studies of vocal ontogeny are indispensable. Rowley and Chapman (1986) showed that galahs (*Eolophus roseicapilla*) cross fostered by Major Mitchell’s cockatoo (*Lophochroa leadbeateri*) produces calls typical to both species, suggesting that calls are socially learned, but also have an innate component, since Major Mitchell’s cockatoo raised by Major Mitchell’s cockatoo parents also appear in mixed flocks but do not produce galah calls. Several studies have shown that parrot chicks go through a babbling phase, where juvenile begging calls and other sounds produced when parents are away from the nest slowly develop into species typical calls (Eggleston et al. 2022; Wein et al. 2021; Berg, Beissinger, and Bradbury 2013; Brittan-Powell et al. 1997). No such study has been done for monk parakeets, but it would be very important to understand when calls are learned, and if they are learned before fledgling, how much they are modified after fledglings gain independence. While we did not study ontogeny directly, our results found no effect of relatedness, either suggesting that individuals modify their contact calls after fledging or that they do not use their parents’ call as a template to begin with. If only a general contact call develops in the nest and birds first learn multiple variants of this call when out of the nest, it could explain the weak signal of spatial distance, since there would be a bias towards learning calls from birds nesting in close proximity. This effect would be weakened once the juveniles disperse. Limited dispersal and relatively high philopatry in monk parakeets in Parc de la Ciutadella would still make it likely for juveniles to nest closer to their parents (Dawson Pell, Senar, D. Franks, et al. 2021), leaving a weak signal of proximity.

Our result, that individuals living in larger nest trees have more diverse repertoires, supports the predictions ofthe social complexity hypothesis– that individuals in larger groups need to communicate more diverse messages (Freeberg 2006). Likewise and more generally, this result supports the predictions made by the primary ratchet model, that larger groups maintain a larger diversity of behaviours (McElreath et al. 2018). We also found that contact call diversity was high across individuals, suggesting that variation in contact call diversity exists within rather than between individuals. The predictions from the social complexity hypothesis are based on production, with individuals producing more variable calls in larger groups because they need to convey more messages (Freeberg 2006). The predictions from the primary ratchet model, on the other hand, are based on acquisition, with individuals acquiring more diverse calls in larger groups, because these larger groups can maintain a more diverse population-level repertoire (McElreath et al. 2018). It should also be pointed out that the predictions from the primary ratched model stem from a mathematical model and have broader implications for any socially learned behaviour. By tracking individuals over longer time spans, it might be possible to ascertain the directionality and separate the predictions from the primary ratched model and social complexity hypothesis. In particular, if individuals disperse to a new nesting location, their vocal diversity and repertoire size should shift if their new nesting tree is different in size. Alternatively, if vocal complexity is related to accumulative experience, dispersal should lead to no change or an increase in diversity and repertoire size.

Previous research has found extensive support for the coevolution of social and vocal complexity across species (Peckre, Kappeler, and Fichtel 2019). There is also an extensive history of work focusing on the drivers of populationlevel differences in vocalisations, for example identifying song cultures and cultural evolution in song (Baker and Jenkins 1987; Aplin 2019). However, studies on the drivers of individual-level differences in vocal complexity are rare. Here, we demonstrate a social influence on call content, diversity and repertoire diversity in a vocal-learning parrot, exhibiting how fine-scale variation in social structure can influence expressed vocal complexity. We further identify that this influence is related to the nesting organisations. Leighton (2017) showed that cooperative breeding birds have a greater diversity of contact and alarm calls. Our results extend this to intra-species variation. Future research could extend this further to examine more species with variable breeding systems, as well as use more detailed sampling to identify the fine-scale social dynamics driving expressed vocal variation.

## Data availability

Data and code are available on GitHub (https://github.com/simeonqs/The_effect_of_social_structure_on_vocal_flexibility_in_monk_parakeets). Some raw data cannot be shared, because it has not been published by our collaborators. The cleaned version is included, so that all results can be reproduced. Some data files are too large for GitHub, these will be included in a permanent Zenodo repository upon acceptance.

## Competing interests

All authors declare they have no competing interests.

## Ethics

The ringing and taking of blood samples was done with special permission EPI 7/2015 (01529/1498/ 2015) from Direcció General del Medi Natural i Biodiversitat, Generalitat de Catalunya, following Catalan regional ethical guidelines for the handling of birds. JCS received authorization (001501-0402.2009) for the handling for research purposes from Servei de Protecció de la Fauna, Flora i Animal de Companyia, according to Decree 214/1997/30.07.

## Funding

SQS received funding from the International Max Planck Research School for Quantitative Behaviour, Ecology and Evolution. LMA was funded by a Max Planck Research Group Leader Fellowship, and is currently supported by the Swiss State Secretariat for Education, Research and Innovation (SERI) under contract number MB22.00056. SQS and LMA were supported by the Centre for the Advanced Study of Collective Behaviour (CASCB), funded by the Deutsche Forschungsgemeinschaft (DFG) under Germany’s Excellence Strategy (EXC 2117-422037984). JCS was supported by a research project from the Ministry of Science and Innovation (CGL-2020 PID2020-114907GB-C21).

### Acknowledgements

We would like to thank Zoo Barcelona and Josep M. Alonso Farré for access to the zoo grounds. We would like to thank Andrés Manzanilla, Gustavo Alarcón-Nieto, Mireia Fuertes Clavero, Alba Ortega-Segalerva and José G. Carrillo-Ortiz for all their support during fieldwork. Finally, we would like to thank Francesca

S. E. Dawson Pell, Daniel Redhead, Mirjam Knö rnschild and Elizabeth Hobson for their advice during the early stages of this project.

## Supplemental materials

### Supplemental methods

#### Relatedness

To establish genetic relatedness, 21 microsatellites that were previously described by Dawson Pell, Senar, D. Franks, et al. (2021) were analysed from the blood samples from 100 individuals. We prioritised analysing samples from individuals for which we had contact calls recorded in 2021, the year for which we also had the best data on social associations. Since samples were stored in ethanol, an extra washing step with phosphase buffered saline was included before DNA extraction. For the DNA extraction PrepManTM Ultra Sample Preparation Reagent (ThermoFisher Scientific) was used according to the manufacturer’s instructions with 5*µ*l of washed blood sample. For the polymerase chain reaction (PCR) the GeneAmp 9700 Thermal Cycler was used with 10 *µ*l reaction volumes containing 1X NH4 buffer, 2 mM MgCl2, 0.2 *µ*M each dNTPs, 0.3 *µ*M of each primer (Dawson Pell, Hatchwell, Ortega-Segalerva, Dawson, et al. 2020), 0.2 U of AmpliTaq GoldTM and 2 *µ*l of diluted DNA 1:4 in molecular water. The thermocycling profile corresponds to a Touchdown polymerase chain reaction (PCR) protocol and the products were separated on the ABI 3500 DNA Analyzer. The full procedure was performed by *Vetgenomics*.

Three samples were analysed twice to ensure all alleles were correctly scored. The resulting allele scores (excluding the reanalysed samples) were analysed in the R package *related* (Pew et al. 2015). We used the function *coancestry* with the settings *wang = 1, trioml*.*num*.*reference = 98*, which returns the pairwise relatedness scores based on Wang (2002). These scores were then normalised between zero and one and used as predictor variables in the contact call similarity analysis.

#### Foraging associations and interactions

We were interested in constructing a social network that captured the full range of social contexts. Monk parakeets spend a great deal of time foraging in fission-fusion flocks. To quantify which conspecifics any individual spent time with, we walked random routes through the park, where the starting point and route were randomised for each session. Data was collected between 27.10.20 - 19.11.20 and 31.10.21 - 30.11.21 (55 days total) with at least six hours of data collection between sunrise and sunset each day. Breaks were scheduled randomly and data collection was only aborted (due to rain or events taking place in the park) for four days. When we encountered a single individual or group foraging or perched in any location other than a nesting tree we recorded the presence of each individual. Unmarked individuals were also counted and recorded, but later excluded from analysis. If birds were being actively provisioned by people or foraging on bread or rice we did not include the recording, because such aggregations might not represent active social association choices. After the group scan was concluded, we then conducted ad libitum observations of displacements and aggressions, allopreening or close tolerance until all individuals had left or 20 minutes had passed. Displacements were defined as any event where one individual showed aggressive behaviour, after which the conspecific moved away. Aggression was defined as pecking, running at or stretching the neck at a conspecific. Close tolerance was defined as two individuals allopreening, touching or beeing oriented towards each other at a distance less than one body length (ca. 10 cm), without any apparent aggression.

### 0.01 Contact call similarity models

#### Foraging network

To model the total effect of the foraging network on vocal similarity we included the nest network, the mate network and the nest cluster network, because they formed forks (see Table 2 for definitions of terms). We also included relatedness, because the nest cluster network was a collider. Since our estimates of acoustic distance between individuals was more certain for individuals for which we had recorded more contact calls, we sampled the *true acoustic distance* from a distribution that took into account the standard deviation in the posterior distribution of the modeled pairwise acoustic distance (*α*_dyad_). Such a construction leads data points with less uncertainty to weigh more. The complete model had the following structure:

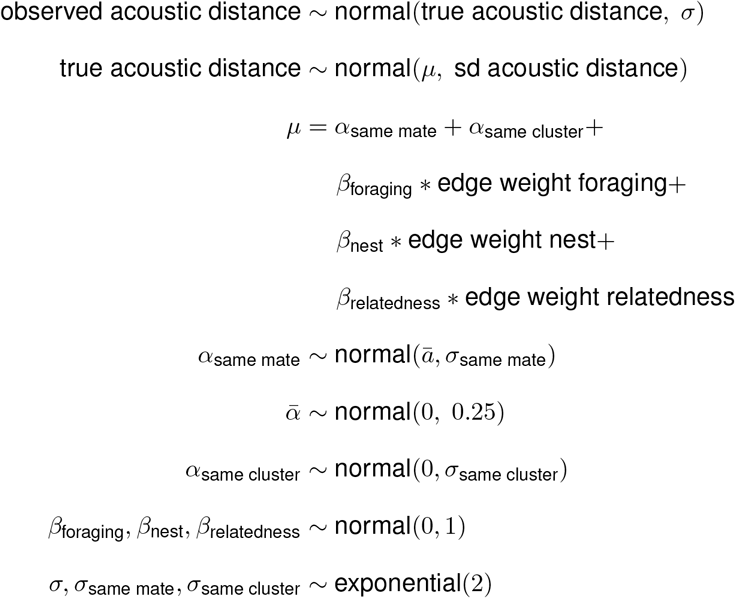

To model the direct effect of the foraging network on vocal similarity we also included the tolerance network, because it formed an indirect path. The model had the following structure:

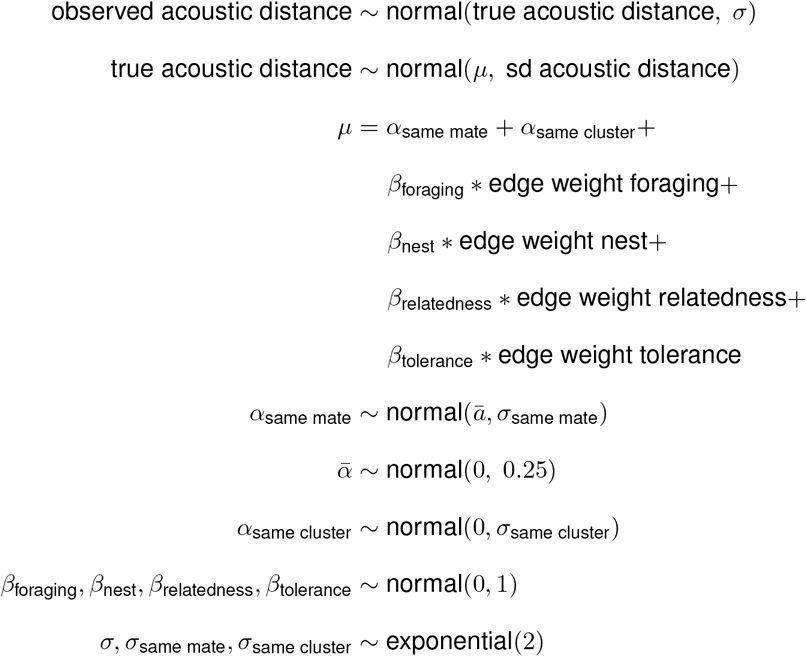

#### Mate network

To model the total effect of the mate network on vocal similarity we only included relatedness, because it formed a fork:

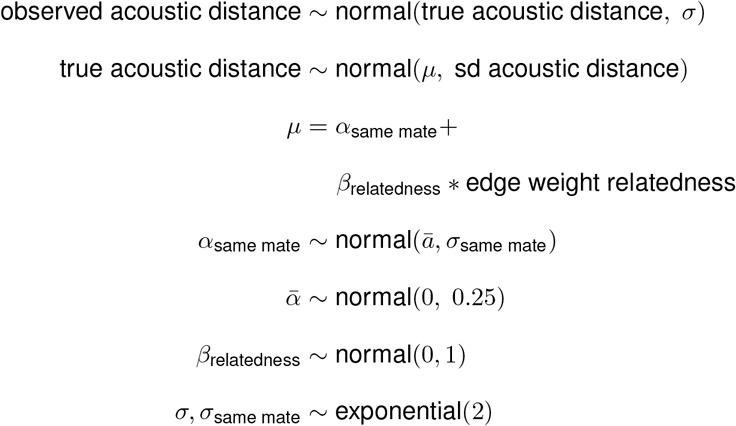

To model the direct effect of the mate network on vocal similarity we also included the foraging and nest networks, which formed indirect paths and the nesting cluster network, because the foraging network was a collider. The model structure was the same as the model for the direct effect of the foraging network.

#### Nest network

To model the total effect of the nest network on vocal similarity we included the nesting cluster network, because it formed a fork. We also included the mate network and relatedness, because the nesting cluster network formed a collider for both these variables. The model structure was as follows:

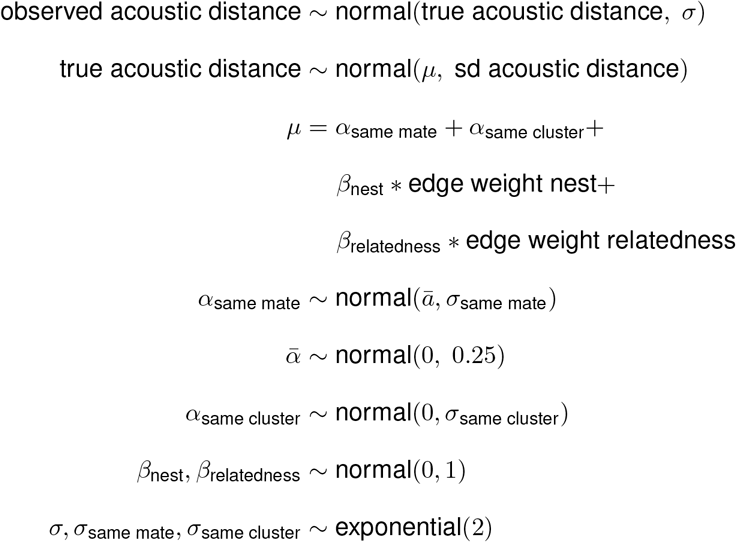

To model the direct effect of the nest network on vocal similarity we additionally included the foraging network, because if was an indirect path. The model structure was the same as the model for the total effect of the foraging network.

#### Nesting network

To model the total effect of the nesting network on vocal similarity we included the mate network and relatedness, because they formed a fork. The model structure was the same as the model for the direct effect of relatedness. To model the direct effect of the nesting network on vocal similarity we also included the foraging, nest and tolerance networks, because they formed indirect paths. The model structure was the same the model for the direct effect of the foraging network.

#### Aggression network

To model the total effect of the aggression network on vocal similarity we included the nesting cluster network, the mate network and relatedness, because they formed forks. The model structure was as follows:

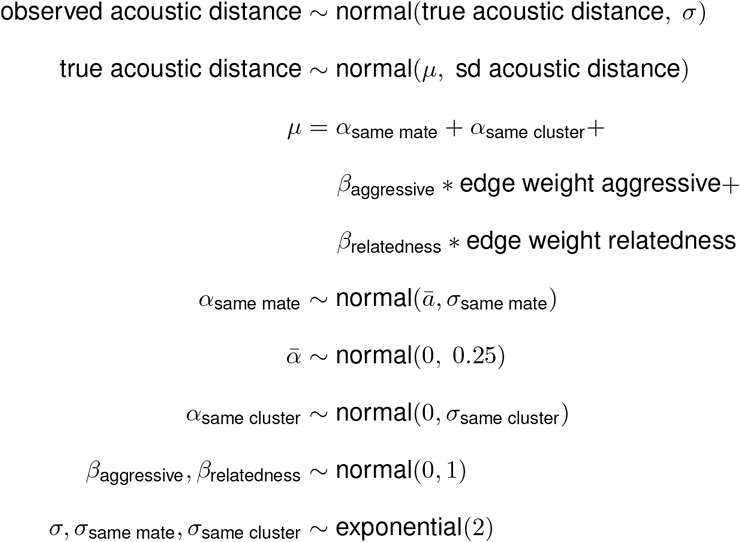

The model for the direct effect of the aggression network on vocal similarity was the same, because there were no indirect paths.

#### Tolerance network

To model the total effect of the tolerance network on vocal similarity we included the foraging, mate and nesting cluster networks, because they were colliders. We also included the nest network and relatedness, because the mate and foraging networks were colliders. The model was the same as the model for the direct effect of the foraging network.

The model for the direct effect of the tolerance network on vocal similarity the model was the same, because there were no indirect paths.

#### Relatedness

To model the total effect of genetic relatedness on vocal similarity we only included relatedness, because there are no arrows entering relatedness. The complete model had the following structure:

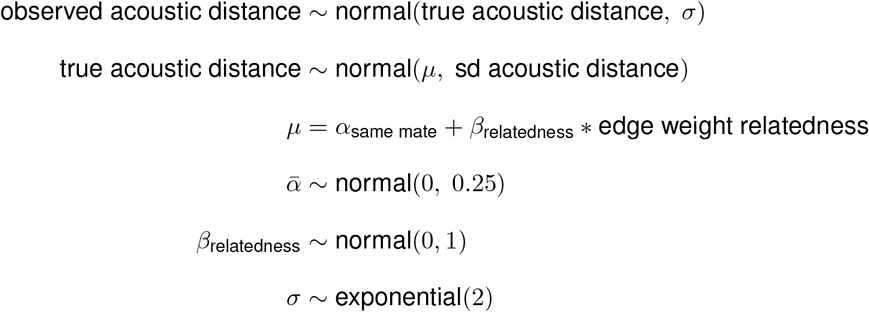

To model the direct effect of the relatedness on vocal similarity we included the mate and nest cluster networks, because they formed indirect paths.

The complete model had the following structure:

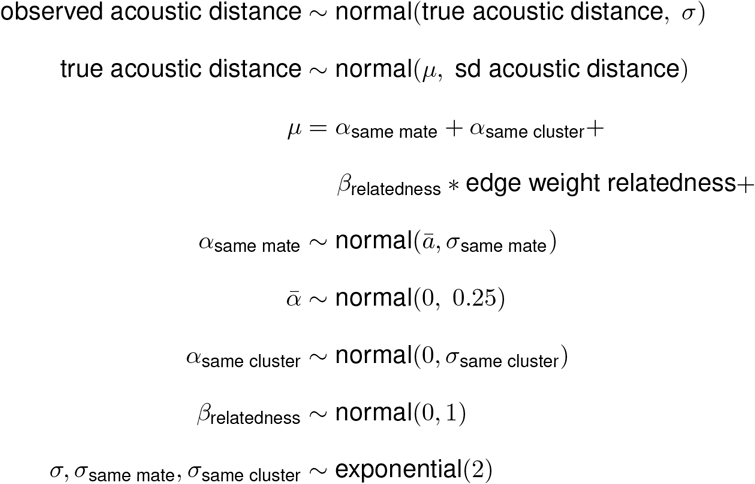

### 00.2 Contact call diversity models

#### Age

To model the total effect of age on individual-level contact call diversity we did not include any covariates since there are no back-door paths into age (see Figure 1b, no arrows entering age). We used the modeled observed variability and standard deviation per individual as response variable in a model with the following structure:

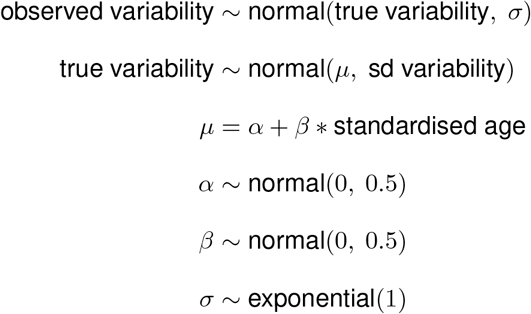

To model the direct effect of age on individual-level contact call diversity we included network position (degree) and chamber size which both are an indirect path. We also included sex, because chamber size is a collider and by including it we opened an indirect path through chamber size and sex to diversity. The model had the following structure:

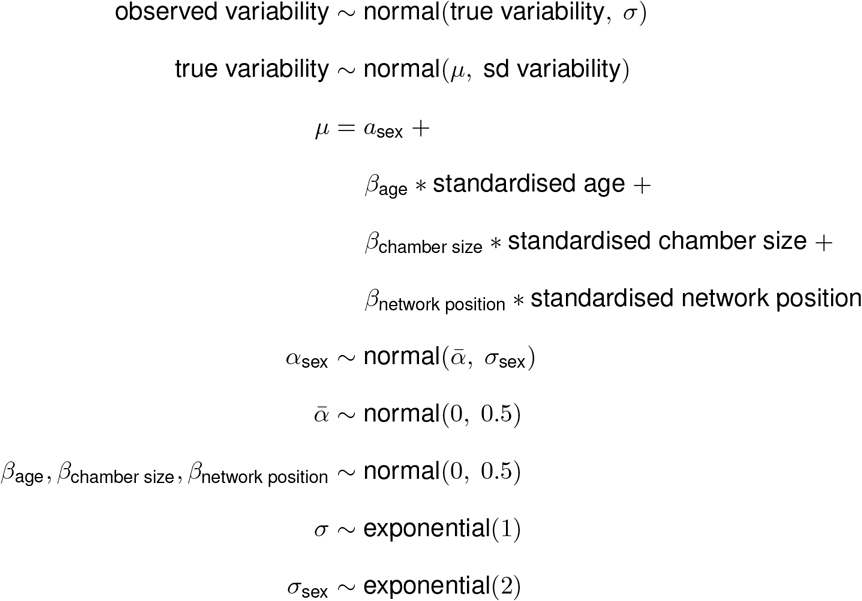

#### Network position

When modelling the effect of network position on individual-level contact call diversity, the model for the total and the direct effect were the same, since there are no arrows going from network position to any other variable than diversity. We did include age, sex, chamber size and tree size as covariates. All these variable were forks. The model had the following structure:

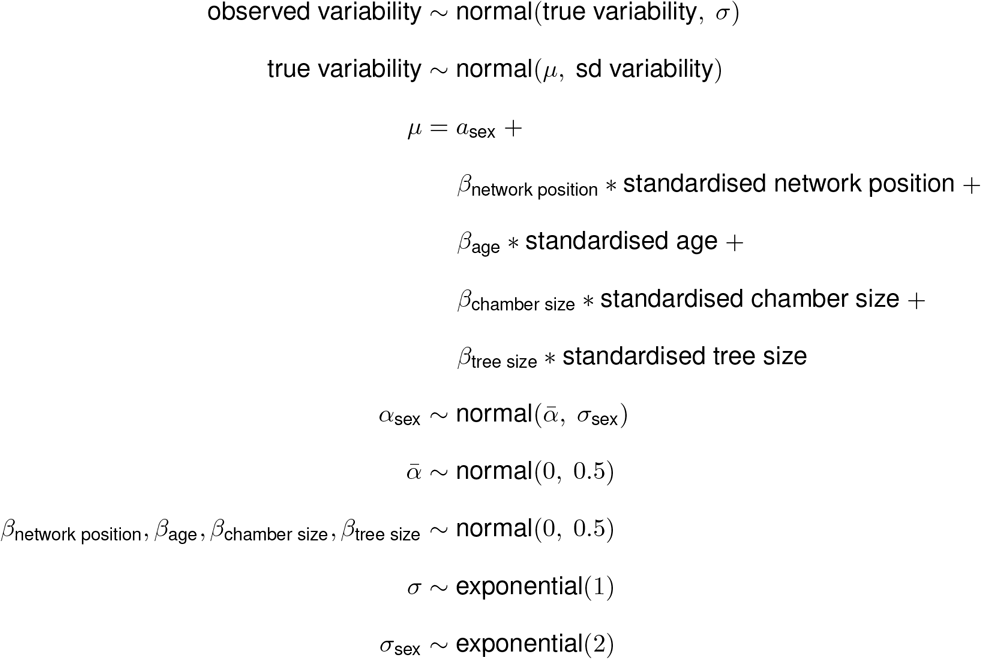

#### Nest chamber size

To model the total effect of nest chamber size on individual-level contact call diversity we included sex and age as covariates. Both were forks. The model had the following structure:

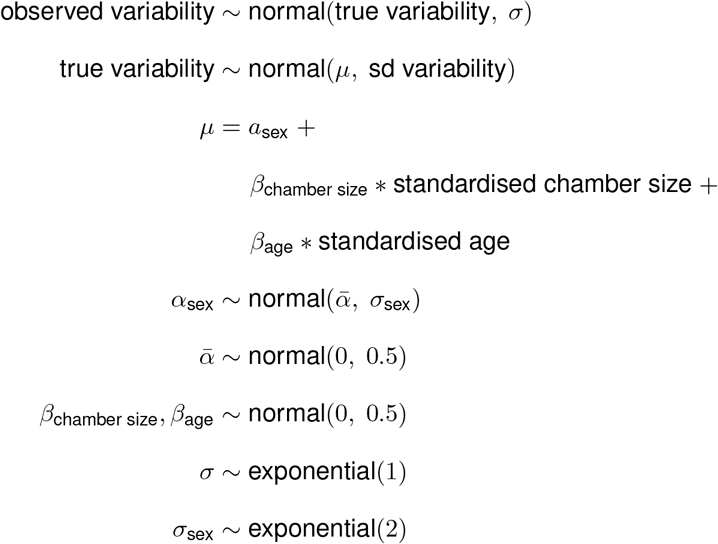

For the direct effect nest chamber size on individual-level contact call diversity we also included tree size, which was an indirect path. The model had the following structure:

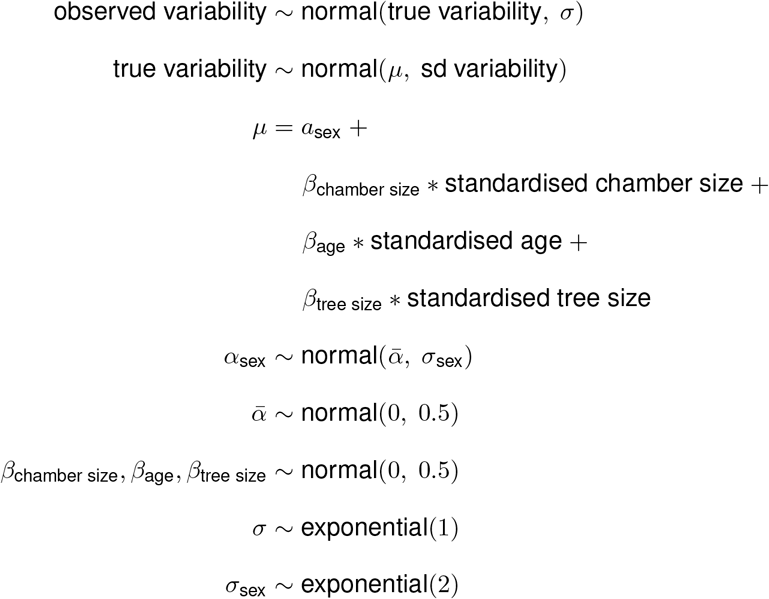

#### Tree size

To model the total effect of tree size on on individual-level contact call diversity we used the same model as for the direct effect of chamber size, because the same covariates needed to be included: chamber size was a fork and a collider that opened paths through age and sex.

To model the direct effect of tree size on individual-level contact call diversity we used the same model as for network position, again because the same covariates needed to be included: chamber size was a fork and a collider that opened paths through age and sex; and network position formed an indirect path.

#### Sex

To model the total effect of sex on on individual-level contact call diversity we did not include any covariates since there are no back-door paths into age. The model had the following structure:

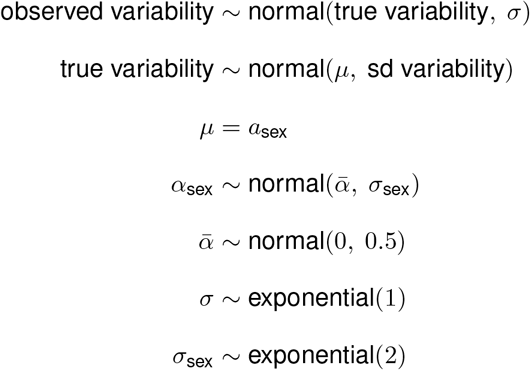

For the direct effect nest chamber size on individual-level contact call diversity we used the same model as for network position, again because the same covariates needed to be included: network position was a fork and a collider that opened paths through age and tree size; and chamber size was an indirect path.

### 0.0.3 Contact call information content models

#### Age

To model the total effect of age on the information content of contact calls in individuals we did not include any covariates since there are no back-door paths into age (see Figure 1c, no arrows entering age). We did include a varying effect for individual, since we included multiple contact calls per individual. The model had the following structure:

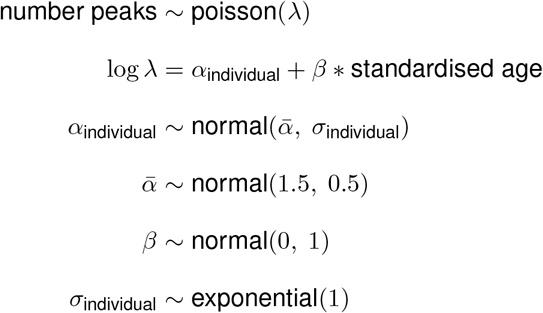

To model the total effect of age on information content of contact calls in individuals we included network position which was an indirect path and tree size, because network position was a collider. The model had the following structure:

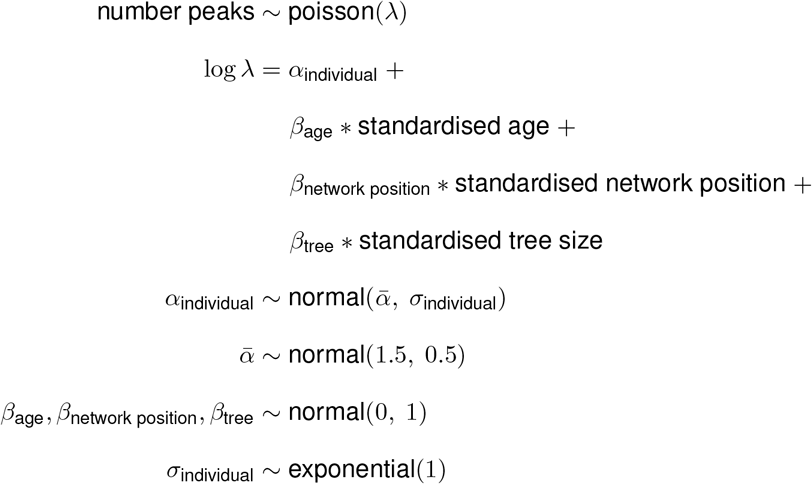

#### Network position

The network position included from the foraging network: degree, eigenvector centrality and betweenness centrality and from the multiplex network it included the degree versatility. We ran separate models for each of these. To model the total effect of network position on information content of contact calls in individuals we included age and tree size, which both formed a fork. The model structure was similar to the model for the direct effect of age.

The direct effect of network position was the same as the total effect.

#### Tree size

To model the total effect of tree size on information content of contact calls in individuals we included chamber size, which formed a fork through network position; age and sex because chamber size was a collider. The model had the following structure:

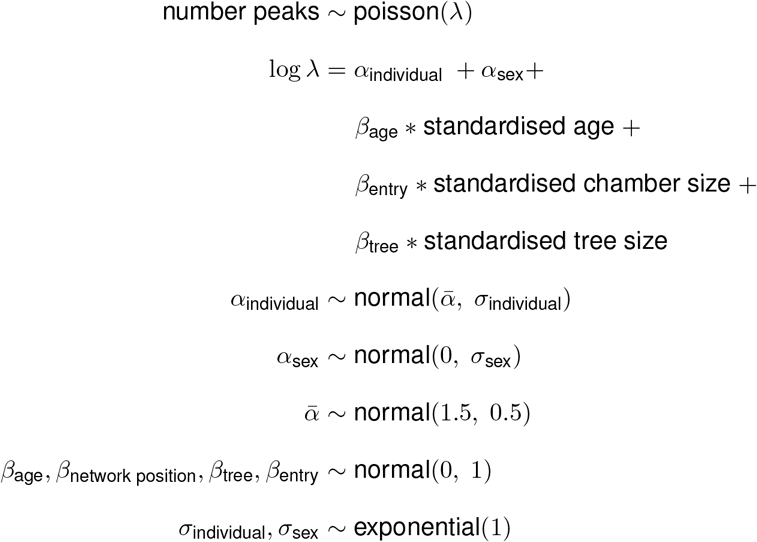

To model the direct effect of tree size on information content of contact calls in individuals we included network position, which was an indirect path, and age, because network position was a collider. The model structure was similar to the model for the direct effect of age.

## Supplemental results

### Model results

### Social networks 2020

### Maps

### Contact call similarity

**Contact call information content**

### Repertoire diversity

## Notes

https://github.com/simeonqs/The_effect_of_social_structure_on_vocal_flexibility_in_monk_parakeets

### Competing Interest Statement

The authors have declared no competing interest.

### Summary of Updates

The manuscript has been revised based on major revision comments from Royal Society Open Science. No changes have been made to the main results or conclusions.

